# Performance evaluation of a new custom, multi-component DNA isolation method optimized for use in shotgun metagenomic sequencing-based aerosol microbiome research

**DOI:** 10.1101/744334

**Authors:** Kari Oline Bøifot, Jostein Gohli, Line Victoria Moen, Marius Dybwad

## Abstract

**Background:** Aerosol microbiome research advances our understanding of bioaerosols, including how airborne microorganisms affect our health and surrounding environment. Traditional microbiological/molecular methods are commonly used to study bioaerosols, but do not allow for generic, unbiased microbiome profiling. Recent studies have adopted shotgun metagenomic sequencing (SMS) to address this issue. However, SMS requires relatively large DNA inputs, which are challenging when studying low biomass air environments, and puts high requirements on air sampling, sample processing and DNA isolation protocols. Previous SMS studies have consequently adopted various mitigation strategies, including long-duration sampling, sample pooling, and whole genome amplification, each associated with some inherent drawbacks/limitations.

**Results:** Here, we demonstrate a new custom, multi-component DNA isolation method optimized for SMS-based aerosol microbiome research. The method achieves improved DNA yields from filter-collected air samples by isolating DNA from the entire filter extract, and ensures unbiased microbiome representation by combining chemical, enzymatic and mechanical lysis. Benchmarking against two state-of-the-art DNA isolation methods was performed with a mock microbial community and real-world subway air samples. All methods demonstrated similar performance regarding DNA yield and community representation with the mock community. However, with subway air samples, the new method obtained drastically improved DNA yields, while SMS revealed that the new method reported higher diversity and gave better taxonomic coverage. The new method involves intermediate filter extract separation into a pellet and supernatant fraction. Using subway air samples, we demonstrate that supernatant inclusion results in improved DNA yields. Furthermore, SMS of pellet and supernatant fractions revealed overall similar taxonomic composition but also identified differences that could bias the microbiome profile, emphasizing the importance of processing the entire filter extract.

**Conclusions:** By demonstrating and benchmarking a new DNA isolation method optimized for SMS-based aerosol microbiome research with both a mock microbial community and real-world air samples, this study contributes to improved selection, harmonization, and standardization of DNA isolation methods. Our findings highlight the importance of ensuring end-to-end sample integrity and using methods with well-defined performance characteristics. Taken together, the demonstrated performance characteristics suggest the new method could be used to improve the quality of SMS-based aerosol microbiome research in low biomass air environments.

## BACKGROUND

The study of bioaerosols is an emerging and expanding research discipline [1], with several important study applications, including surveillance of clinically relevant microbes [2–5], air quality monitoring [6–8] and biodefense [9]. Bioaerosol research has traditionally relied on culture methods; however, few microorganisms grow under standard laboratory conditions, resulting in underrepresentation of the true microbial diversity [10–13]. Although culture methods are still in use, culture-independent methods are now widespread. Due to the low amount of DNA that is typically obtained from air samples, most culture-independent bioaerosol studies to date have used PCR to target either the bacterial 16S rRNA gene [14, 15] or the fungal 18S rRNA gene/internal transcribed spacer (ITS) region, followed by amplicon sequencing [16, 17]. In contrast to the amplicon sequencing approach, shotgun metagenomic sequencing (SMS) allows for generic, unbiased interrogation of microbial diversity in a sample. However, SMS will typically require a higher quality and quantity of DNA for analysis than other molecular methods. SMS has been used to characterize the human microbiome [18] and environmental microbiomes [19, 20], and has recently been implemented in a few aerosol microbiome studies [2, 21, 22].

Although bioaerosols originate from many different sources and are ubiquitous in almost any indoor and outdoor environment, air is still a very low biomass environment compared to e.g. soil, feces and water [23]. The low biomass makes it challenging to obtain sufficient DNA amounts for downstream analyses, especially in the context of SMS [21]. An important first step in recovering sufficient biomass and a representative sample from air involves the use of well-characterized air samplers that are capable of rapid and efficient biomass collection [24]. Filter-based aerosol collection is a commonly used method, and the use of hand-portable, high-volume filter-based air sampling equipment may improve the spatiotemporal resolution in aerosol microbiome research [24, 25]. The post-sampling processing steps are also important since the filter-collected biomass must be transformed into a representative high quality DNA sample with minimal loss. It is therefore essential to use a well-characterized DNA isolation method that is capable of thorough unbiased biomass lysis, sufficient inhibitor removal and sample clean-up, and high efficiency recovery of DNA [25]. In short, the main challenges are typically obtaining sufficient DNA amounts and capturing representative samples that reflect the true diversity of the sampled air environment [2, 22, 25, 26].

With recent advancements in sequencing technology, along with the development of improved strategies for air sampling and sample processing, it should be possible to mitigate the low biomass challenge. Mitigation strategies that have been attempted in the past include long-duration sampling (days to weeks), pooling of multiple air samples, whole genome amplification (WGA) techniques, and modification of commercial DNA isolation kits originally developed for other environmental matrices such as water and soil [2, 21, 27–29]. Increasing the air sampling time is a common strategy to improve the DNA yield, but this approach may not always be practical. For example, in studies where the aim is to address spatiotemporal variability, the need for long-duration air sampling (e.g. days to weeks) exclude the possibility of aerosol microbiome investigations on shorter timescales. Another challenge with increased air sampling time is that long-duration filter collection may compromise the integrity of stress-sensitive microorganisms, e.g. due to desiccation and osmotic shock [27], and thereby cause a potential loss of DNA from organisms that become membrane-compromised, ruptured or lysed during filter extraction and subsequent processing steps prior to DNA isolation. Liquid extraction of aerosol filters often results in sample volumes that are too large to process with most commercial DNA isolation kits. This introduces a need for adopting additional post-extraction filtration or centrifugation steps to reduce the sample volume before DNA isolation, which may result in loss of both intact microorganisms and DNA, and thereby compromise the sample integrity regarding both yield and composition (diversity). Furthermore, long-duration, high-volume air sampling alone does not always translate into successful recovery of sufficient DNA amounts for SMS [2, 21, 28, 29]. This may be due to the use of different downstream sample processing and DNA isolation methods that have not been sufficiently evaluated regarding their specific performance on air samples, and which therefore may deliver suboptimal performance regarding biomass lysis and/or DNA recovery efficiency. Various modifications of existing sample processing and DNA isolation methods have been proposed to improve the DNA yield from filter-collected air samples. Jiang et al. modified the DNeasy (former MO-BIO) PowerSoil Kit by replacing the silica spin column with AMPure XP beads, and introduced sample pre-treatment steps and a secondary filtration step [28]. Yooseph et al. introduced a WGA step to generate sufficient DNA amounts from air samples for SMS [21]. King et al. performed liquid extraction of aerosol filters followed by a secondary filtration step and DNA isolation with the DNeasy PowerWater Kit, and precipitated DNA from the original filtrate before combining the two DNA fractions [2]. Dommergue et al., who also used the DNeasy PowerWater Kit, placed the aerosol filters directly in PowerBead tubes, introduced sample pre-treatment steps, and a centrifugation step to maximize lysate recovery from PowerBead tubes [29]. Recovery of sufficient DNA amounts and preservation of unbiased microbial diversity from air samples is essential to ensure reliable results in SMS-based aerosol microbiome research. Several studies on other sample matrices have looked into how DNA yields can be improved and microbial diversity preserved. Tighe et al. found that using a multi-enzyme cocktail (MetaPolyzyme) that targets bacterial and fungal cell wall components resulted in improved DNA yields [30]. Yuan et al. evaluated different DNA isolation methods for human microbiome samples, and found bead beating and enzymatic lysis to be essential for obtaining an accurate representation of microorganisms in a complex mock community [31]. Abusleme et al. found that bead beating may limit the DNA yield, but also that bead beating was necessary to detect all organisms in a complex mock bacterial community [32]. These observations suggest that biomass lysis based on a combination of chemical, enzymatic and mechanical principles may be useful to minimize microbiome composition (diversity) bias resulting from insufficient biomass lysis during isolation of DNA from complex environmental assemblages.

It is well established that the choice of DNA isolation method should be based on careful consideration of the specific study aims, including type of targeted organisms and environmental matrices [33]. However, substantial uncertainty exists regarding the extent of microbiome composition (diversity) bias that may be introduced by the use of different sample processing and DNA isolation methods, which makes it difficult to reliably compare microbiome results between different studies and environments. Consequently, several attempts have in recent years been made to improve the harmonization and standardization of DNA isolation methods, especially for common sample matrices such as human [31, 34], soil [35], and water [36] samples. Lear et al. recommended DNA isolation kits for different environmental matrices such as soil, plant and animal tissue, and water [37]. The Earth Microbiome Project demonstrated how procedural standardization allows for comparison of microbial diversity in samples from across the globe [35]. Dommergue et al. proposed an air sampling, filter extraction and DNA isolation method where microbial diversity and chemical composition in air can be investigated using existing high-volume particulate matter samplers used for atmospheric pollution monitoring [29]. Nevertheless, despite substantial effort several unresolved issues remain, e.g., the current reliance on long-duration air sampling raises some questions regarding sample integrity and only offers support for low temporal resolution studies since the necessary sampling time may be days or even weeks. Hence, performance benchmarking, harmonization, and standardization of air sampling, sample processing and DNA isolation methods is a topic that warrants further study, and especially in the context of SMS-based aerosol microbiome research, which is a research field still largely in its infancy.

The aim of this study was to demonstrate a new custom, multi-component DNA isolation method optimized for SMS-based aerosol microbiome research and perform a comprehensive performance benchmarking of the new method. The custom, multi-component DNA isolation method was specifically developed to maximize the DNA yield and ensure unbiased biomass lysis from low biomass environmental air samples. The DNA isolation method, hereafter referred to as the “MetaSUB method”, was developed for the MetaSUB Consortium (www.metasub.org) to complement an ongoing global effort to characterize subway and urban environment microbiomes using surface swab samples, by extending the effort to also include air samples. The MetaSUB method was benchmarked against two other state-of-the-art DNA isolation methods: a custom multi-component DNA isolation method developed for use in aerosol microbiome research published by Jiang et al. [28], and the commercial ZymoBIOMICS DNA Microprep Kit commonly used in environmental microbiome studies [38–41]. The performance of the three DNA isolation methods was evaluated using both a mock microbial community and real-world low biomass subway air samples. As part of this study, we also describe an end-to-end high-volume filter-based air sampling, filter processing and DNA isolation method, hereafter referred to as the “end-to-end MetaSUB method”. Since the MetaSUB method, when used as an integrated element of the end-to-end MetaSUB method, involves intermediate separation of the filter extract into a pellet (subjected to additional lysis) and supernatant fraction that is combined before final DNA purification, the relative contribution of the two fractions to the total DNA yield and observed aerosol microbiome profile was also evaluated using subway air samples.

## METHODS

### MetaSUB method

The end-to-end MetaSUB method consists of an integrated air sampling, filter processing and DNA isolation scheme (Figure 1). The method relies on the use of high-volume, battery-operated, hand-portable, electret filter-based air samplers that allow for flexible, user-adjustable sampling time and rapid change of sampling locations, which in turn provides support for high spatiotemporal resolution air (aerosol biomass) sampling campaigns. Following air sampling, the electret microfibrous filter is subjected to a liquid filter extraction procedure, after which the entire filter extract is processed to avoid the need for downstream filtration or centrifugation steps to reduce the sample volume prior to DNA isolation, which may compromise the sample integrity regarding both biomass and DNA yield and composition (diversity).

**Figure 1.**
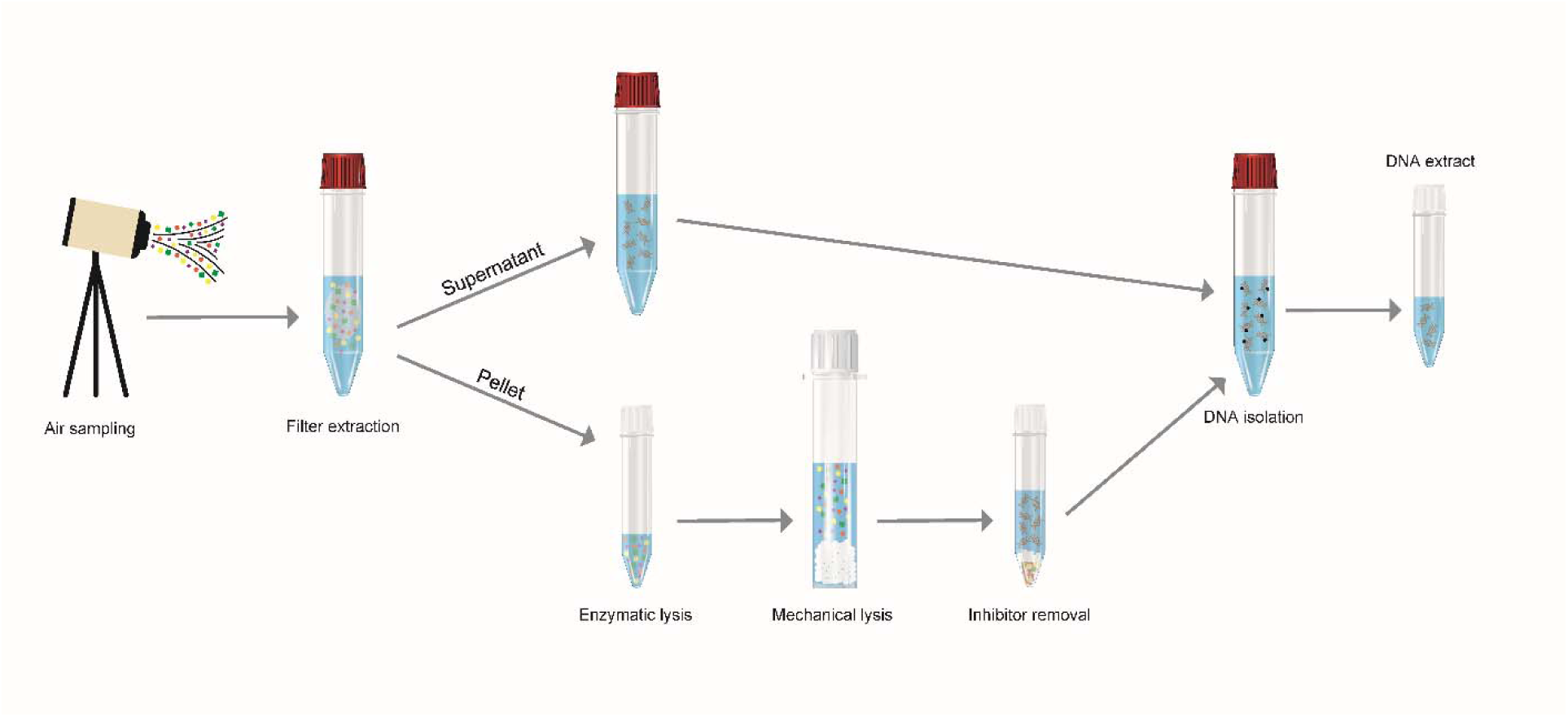
Overview of the end-to-end MetaSUB method. Air samples collected using SASS 3100 high-volume filter-based air samplers (Research International) on SASS 3100 electret microfibrous filters (Research International) are extracted in NucliSENS lysis buffer (BioMérieux) and centrifuged, resulting in intermediate separation of the filter extract into a pellet and supernatant fraction. The pellet is subjected to additional lysis with MetaPolyzyme (Sigma-Aldrich), a multi-enzyme cocktail, followed by bead beating with ZR Bashing Tubes (Zymo Research) filled with PowerSoil Bead Solution (Qiagen) and Solution C1 (Qiagen). Inhibitor removal and sample clean-up is performed with Solution C2 and C3 (Qiagen). The supernatant and pellet fractions are recombined and DNA purification performed according to the manual protocol of the NucliSENS Magnetic Extraction Reagents kit (BioMérieux).

### Bioaerosol collection

Air (aerosol biomass) samples were collected with SASS3100 (Research International, Monroe, WA, USA), a high-volume electret microfibrous filter-based air sampler. The air sampler was powered by UBI-2590 lithium-ion rechargeable batteries (Ultralife batteries, NY, USA), operated at a flowrate of 265 liters of air per minute (LPM), and mounted on a tripod (∼1.5 meters above ground) with the inlet facing downward (45°) to avoid direct deposition of large particles. After sampling, the electret filters were stored in 50 ml polypropylene tubes at −80 °C until further processing.

### Filter extraction

Liquid extraction of filter-collected aerosol biomass from the electret filters was performed by removing the filters from their housing and transferring them into 50 ml polypropylene tubes pre-loaded with 10 ml NucliSENS Lysis Buffer (BioMérieux, Marcy-l’Étoile, France). The sample tube was vortexed at maximum speed for 20 seconds before the filter was transferred into a 10 ml syringe with sterile forceps to extract residual liquid back into the sample tube before discarding the filter. The sample tube was centrifuged (7000 x g, 30 minutes) and the supernatant transferred to a new 50 ml polypropylene tube (referred to as filter extract supernatant).

### DNA isolation

The pellet from the sample tube (referred to as filter extract pellet) was transferred to a polypropylene microcentrifuge tube with 1 ml PBS (pH 7.5, Sigma-Aldrich, St. Louis, MO, USA) and centrifuged (17 000 x g, 5 minutes). The resulting supernatant was carefully removed and combined with the filter extract supernatant. The pellet was dissolved in 150 µl PBS (pH 7.5). MetaPolyzyme (Sigma-Aldrich), a multi-enzyme cocktail, was prepared by dissolving the enzyme powder in 1 ml PBS (pH 7.5), and 10 µl MetaPolyzyme (5 mg/ml) and 5 µl sodium azide (0.1 M, Sigma-Aldrich) was added to the dissolved pellet sample. Enzymatic digestion was performed at 35°C for 1 hour in a Thermomixer (Eppendorf, Hamburg, Germany) at 1400 rpm. Subsequently, the sample was transferred to ZR BashingBead Lysis Tubes (0.1/0.5 mm beads, Zymo Research, Irvine, CA, USA) prefilled with 550 µl PowerSoil Bead Solution (Qiagen, Hilden, Germany) and 60 µl PowerSoil Solution C1 (Qiagen). Bead tubes were subjected to bead beating (17 000 x g, 3 minutes) in a Mini Bead Beater-8 (BioSpec Products, Bartlesville, OK, USA). Bead tubes were centrifuged (13 000 x g, 2 minutes) and the supernatant treated with Solution C2 and C3 according to the DNeasy PowerSoil protocol (Qiagen). The resulting supernatant was combined with the original filter extract supernatant before DNA purification. DNA was purified according to the manual protocol of the NucliSENS Magnetic Extraction Reagents kit (BioMérieux) with two modifications; magnetic silica suspension volume was increased to 90 µl and incubation time was increased to 20 minutes. DNA samples were stored at −80°C until further processing.

### DNA isolation method described by Jiang et al. (Jiang method)

The custom, multi-component DNA isolation method (protocol steps 13-24) for air samples published by Jiang et al. [28] is based on the DNeasy PowerSoil Kit and AMPure XP magnetic bead separation. Jiang et al. introduced an incubation step in water bath (65°C) before bead vortexing, and found that magnetic bead capture recovered more DNA than standard PowerSoil spin columns. The DNA isolation method (protocol steps 13-24) published by Jiang et al. (hereafter referred to as “Jiang”) was used in this study with some minor modifications. Briefly, all samples were pretreated with MetaPolyzyme (as described for the MetaSUB method), before transfer to PowerBead tubes and continuation of DNA isolation according to the Jiang protocol.

### ZymoBIOMICS DNA Microprep Kit (Zymobiomics method)

DNA isolation was performed according to the ZymoBIOMICS DNA Microprep Kit (Zymo Research) protocol (hereafter referred to as “Zymobiomics”) with some minor modifications. Briefly, all samples were pretreated with MetaPolyzyme (as described for the MetaSUB method) and bead beating was performed in a Mini Bead Beater-8 (BioSpec Products) for 3 minutes.

### Performance evaluation using mock microbial community

The MetaSUB method was compared to the Jiang and Zymobiomics methods using a mock microbial community with a defined quantity and composition. The ZymoBIOMICS Microbial Community Standard (Zymo Research) contains ten microorganisms, eight bacteria (five Gram-positives and three Gram-negatives) and two yeasts. For each sample, the mock community (10 µl), corresponding to a theoretical total DNA content of approximately 267 ng, was added to 140 µl PBS (pH 7.5) and treated with MetaPolyzyme (as described for the MetaSUB method) before DNA isolation according to the three DNA isolation methods. Total DNA and 16S rRNA gene copy yields were measured for four sample pairs processed with MetaSUB (N=4) and Jiang (N=4) and six sample pairs processed with MetaSUB (N=6) and Zymobiomics (N=6). The within-sample differences in total DNA and 16S rRNA gene copy yields were evaluated with one-sample t-tests (H_0_: difference=0). All statistical analyses were performed in R (version3.4.3, www.R-project.org). A subset of the mock community samples were subjected to SMS (N=12): MetaSUB (N=4), Jiang (N=4), and Zymobiomics (N=4).

### Performance evaluation using subway air samples

The MetaSUB method was compared to the Jiang and Zymobiomics methods using subway air samples. Only the DNA isolation part of the end-to-end MetaSUB method was evaluated since the air sampling and filter-processing steps were used to collect and process subway air samples to generate equal aliquots of aerosol biomass for paired difference comparisons. An overview of the common sample processing steps and the three evaluated DNA isolation methods is given in Table 1. Air samples were collected for 1 hour, corresponding to ∼16 m^3^ of air sampled (60 minutes sampling at 265 LPM), during daytime hours at subway stations (Tøyen, Grønland, Stortinget, Nationaltheateret and Majorstuen) in Oslo, Norway, in the period between October 2017 and May 2018. The filter-collected samples were extracted in 10 ml NucliSENS lysis buffer and split into two equal filter extract aliquots. The aliquots were centrifuged (7000 x g, 30 minutes) and only the pellet fractions were used for the comparison of DNA isolation methods. The supernatant fractions were subjected to DNA isolation separately (as described below) and used to investigate the distribution of DNA in the intermediate pellet and supernatant fractions of the MetaSUB method. For the DNA isolation method comparison, 24 air samples were split and the pellets processed with either MetaSUB (N=10) and Jiang (N=10) or MetaSUB (N=14) and Zymobiomics (N=14), to enable within-sample comparisons between the MetaSUB method and the two other methods. Since the supernatant fraction was not included in the MetaSUB method for the DNA isolation method comparison, 10 ml of fresh NucliSENS lysis buffer was used. Negative controls (reagents) were included for each DNA isolation method. Total DNA and 16S rRNA gene copy yields were examined and within-sample differences were evaluated with one-sample t-tests (H_0_: difference=0). All statistical analyses were performed in R (version3.4.3, www.R-project.org). A subset of the subway air samples (N=6) that had been split into two equal aliquots and processed with the three DNA isolation methods were subjected to SMS (N=12): MetaSUB (N=3) v. Jiang (N=3) and MetaSUB (N=3) v. Zymobiomics (N=3). A negative control (reagents) for each DNA isolation method was also subjected to SMS (N=3).

**Table 1.** Overview of the three DNA isolation methods evaluated in this work.

| Method | MetaSUB | Jiang | Zymobiomics |
| --- | --- | --- | --- |
| **Common processing steps**<br>(used to generate equal aerosol biomass aliquots for paired difference comparison) |  |  |  |
| Filter extraction<br>(filter-to-liquid) | NucliSENS lysis buffer |  |  |
| Lysis (enzymatic) | MetaPolyzyme multi-enzyme cocktail |  |  |
| **Method-specific processing steps**<br>(used for paired difference comparison on equal aerosol biomass aliquots) |  |  |  |
| Lysis (mechanical) | ZR BashingBead Tubes with PowerSoil Bead Solution and Solution C1. Bead beating for 3 min | PowerSoil Bead Tubes with PowerSoil Bead Solution and Solution C1 incubated at 65°C for 15 min. Bead vortexing for 15 min | ZR BashingBead Tubes with Zymobiomics lysis solution. Bead beating for 3 min |
| Inhibitor removal and sample clean-up | PowerSoil Solution C2 and C3 | PowerSoil Solution C2 and C3 | Zymo-Spin IV and Zymo-Spin IV-μHRC Columns |
| DNA purification | NucliSENS magnetic beads | AMPure XP magnetic beads | Zymo-Spin IC-Z Column |

### DNA distribution in intermediate pellet/supernatant fractions of the MetaSUB method

The filter extraction procedure of the MetaSUB method generates two intermediate fractions (pellet and supernatant) that are usually recombined before the final DNA purification (Figure 1). Differences in total DNA and 16S rRNA gene copy yields between pellet (N=24) and supernatant (N=24) fractions were therefore investigated. DNA was isolated from the supernatant fractions from subway air samples (described above) with the NucliSENS Magnetic Extraction Reagents kit as described for the MetaSUB method. Furthermore, to identify potential differences in DNA composition (diversity) between the pellet and supernatant fractions, DNA isolated with the MetaSUB method from six paired pellet and supernatant fractions (N=12) and one negative control (reagents; N=1) was subjected to SMS.

### Quantification of total DNA and 16S rRNA gene copies

Total DNA was quantified with Qubit dsDNA HS assays (Life Technologies, Carlsbad, CA, USA) on a Qubit 3.0 Fluorimeter (Life Technologies). Bacterial 16S rRNA gene copies were determined with a 16S rRNA gene qPCR assay performed according to Liu et al. [42] on a LightCycler 480 instrument (Roche Diagnostics, Oslo, Norway). Serial dilutions of *Escherichia coli* DNA (seven 16S rRNA gene copies per genome) were used to generate a standard curve.

### Shotgun metagenomic sequencing (SMS)

DNA isolated from mock community samples were subjected to SMS (150 bp paired-end) multiplexed on a MiSeq (∼24-30 M paired-end reads, Illumina, San Diego, CA, USA). Library preparation was done with the Nextera DNA Flex kit (Illumina) according to the recommended protocol. DNA isolated from subway air samples were subjected to SMS (150 bp paired-end) multiplexed on one lane (∼80-130M paired-end reads) on a HiSeq 3000 (Illumina). Library preparation was done with the ThruPLEX DNA-Seq kit (Takara Bio, Mountain View, CA, USA) according to the recommended protocol and 18 amplification cycles. Raw sequence reads were demultiplexed, quality trimmed (Trim Galore, v0.4.3; ≥Q20, ≥50 bp) and underwent adapter removal (Cutadapt, v1.16), before analysis on the One Codex platform with default settings [43]. One Codex taxonomic feature tables were imported into R and analyzed in the phyloseq package [44].

All sequence reads not taxonomically assigned to the species level were removed from the 12 mock community samples. Since the aim was to gauge the relative contribution of the ten bacterial and fungal species in the mock community across the three DNA isolation methods, non-target features were binned as “other”. The comparison was made by plotting normalized abundances across all 12 samples.

For the six subway air samples that were split into equal aliquots and processed with the three DNA isolation methods, MetaSUB (N=3) v. Jiang (N=3) and MetaSUB (N=3) v. Zymobiomics (N=3), all taxonomic features not assigned to the genus or species level, along with human reads, were removed. Prevalent features reported in the negative control samples (>1% of within-sample reads, four in total, accounting for 94.5% of all reads in the negative controls) were stripped from the entire dataset before removing the negative controls. The cleaned samples varied in the number of assigned reads, ranging from 1 160 976 to 5 530 138. After examining the effect of rarefication on the α-diversity measures “Observed”, “Shannon”, and “Simpson” (Figure S1), all samples were rarified to the lowest common depth (1 160 976).

The six paired pellet and supernatant fractions from subway air samples processed with the MetaSUB method underwent the same procedure: removing features not assigned to the genus or species level, along with human reads, and prevalent features in the negative control (12 features, accounting for 99.3% of all reads in the negative control). The effect of rarefication was evaluated (Figure S2), and all samples were rarified to the lowest common depth (453 218).

The cleaned SMS datasets were divided into six groups corresponding to the three comparisons (MetaSUB v. Jiang, MetaSUB v. Zymobiomics, and MetaSUB pellet v. supernatant) before summarizing the top phyla, families, genera and species within each group. Taxonomic features with species-level assignment were extracted for analyses of within-sample diversity (α-diversity: “Observed”, “Shannon”, “Simpson”), where relevant groups were compared by fitting linear models. All features (read counts) were conglomerated to the genus level for analyses of among sample differences (β diversity); Bray Curtis distances were ordinated with PCoA and analyzed with MetaSUB/Jiang, MetaSUB/Zymobiomics, and pellet/supernatant, as predictors in separate PERMANOVA tests. Distance estimation and PERMANOVA was performed with vegan (v.2.6.0, https://github.com/vegandevs/vegan/). Sample clustering was visualized with PCoA ordination. MegaBLAST analysis of forward reads against the NCBI non-redundant nucleotide database, followed by taxonomic binning using the native lowest common ancestor (LCA) algorithm in MEGAN6 [45], was used to perform a cross-kingdom analysis on the pellet/supernatant samples. Lastly, random forest classification models were performed, using 10 001 trees, with MetaSUB/Jiang, MetaSUB/Zymobiomics, and pellet/supernatant, as response variable and One Codex (species-level) taxonomic features as predictor variables. Separate tests using 501 trees and 1000 permutations were performed to evaluate statistical significance. The random forest models were built using randomForest [46].

### Accession numbers

The sequence data has been deposited in the NCBI Sequence Read Archive under Bioproject ID# PRJNA542423 (https://www.ncbi.nlm.nih.gov/bioproject/PRJNA542423).

## RESULTS

### Performance evaluation using mock microbial community

The total DNA and 16S rRNA gene copy yields from mock community samples showed no significant differences between the MetaSUB method and the other two methods (Figure 2; Table 2A). However, the MetaSUB method obtained a higher 16S rRNA gene copy yield than Jiang with borderline significance (*P* = 0.055; Figure 2; Table 2A). The 12 mock community samples that were subjected to SMS showed similar distributions of all ten microbial species in the mock community across the three methods, with MetaSUB and Zymobiomics being nearly identical (Figure 3).

**Figure 2.**
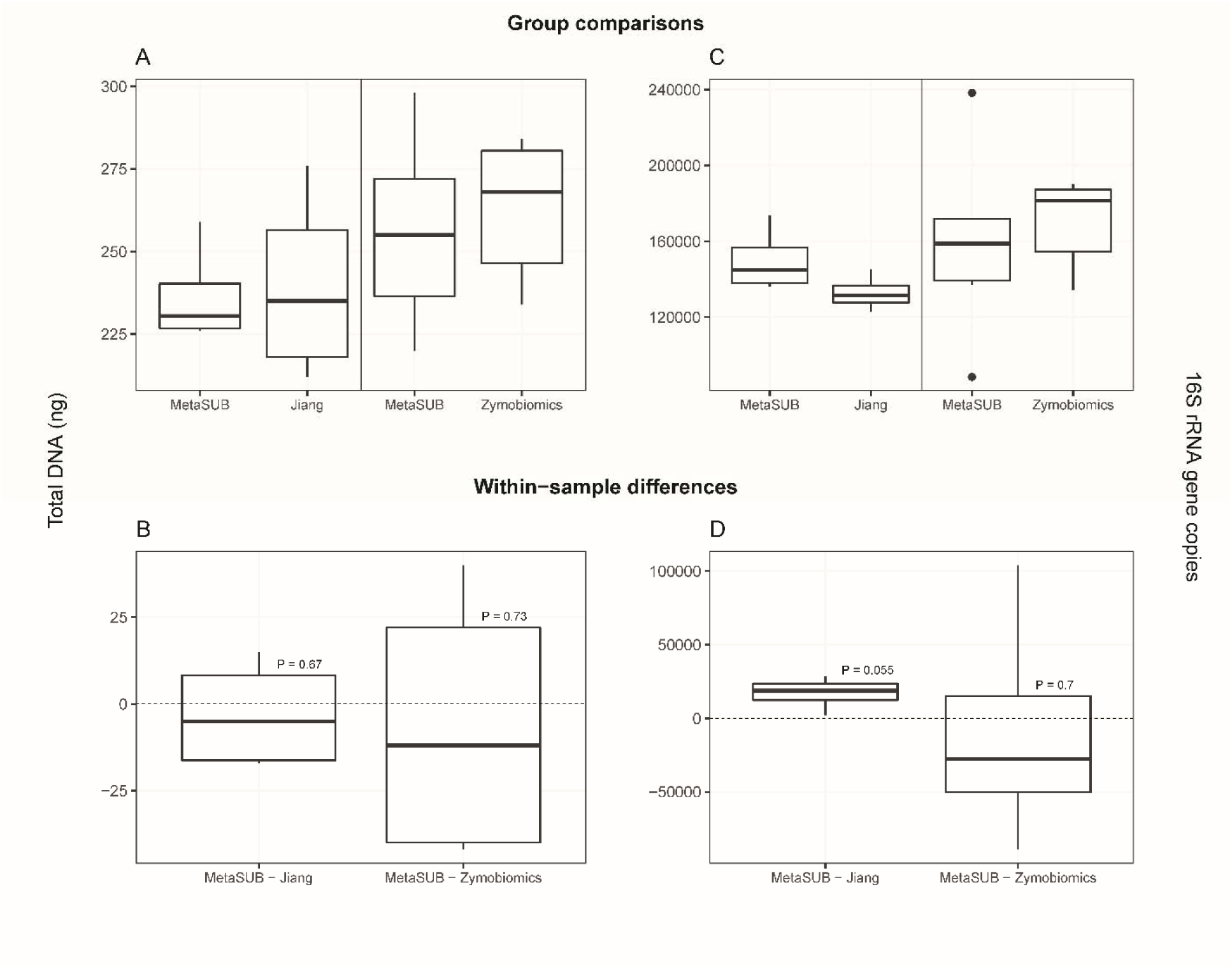
Benchmarking results for MetaSUB, Jiang, and Zymobiomics on mock microbial community samples. One sample t-tests were performed on within-sample differences (B, D) of total DNA yield (A), and 16S rRNA gene copy yield (C) for MetaSUB (N=4) and Jiang (N=4), and MetaSUB (N=6) and Zymobiomics (N=6).

**Figure 3.**
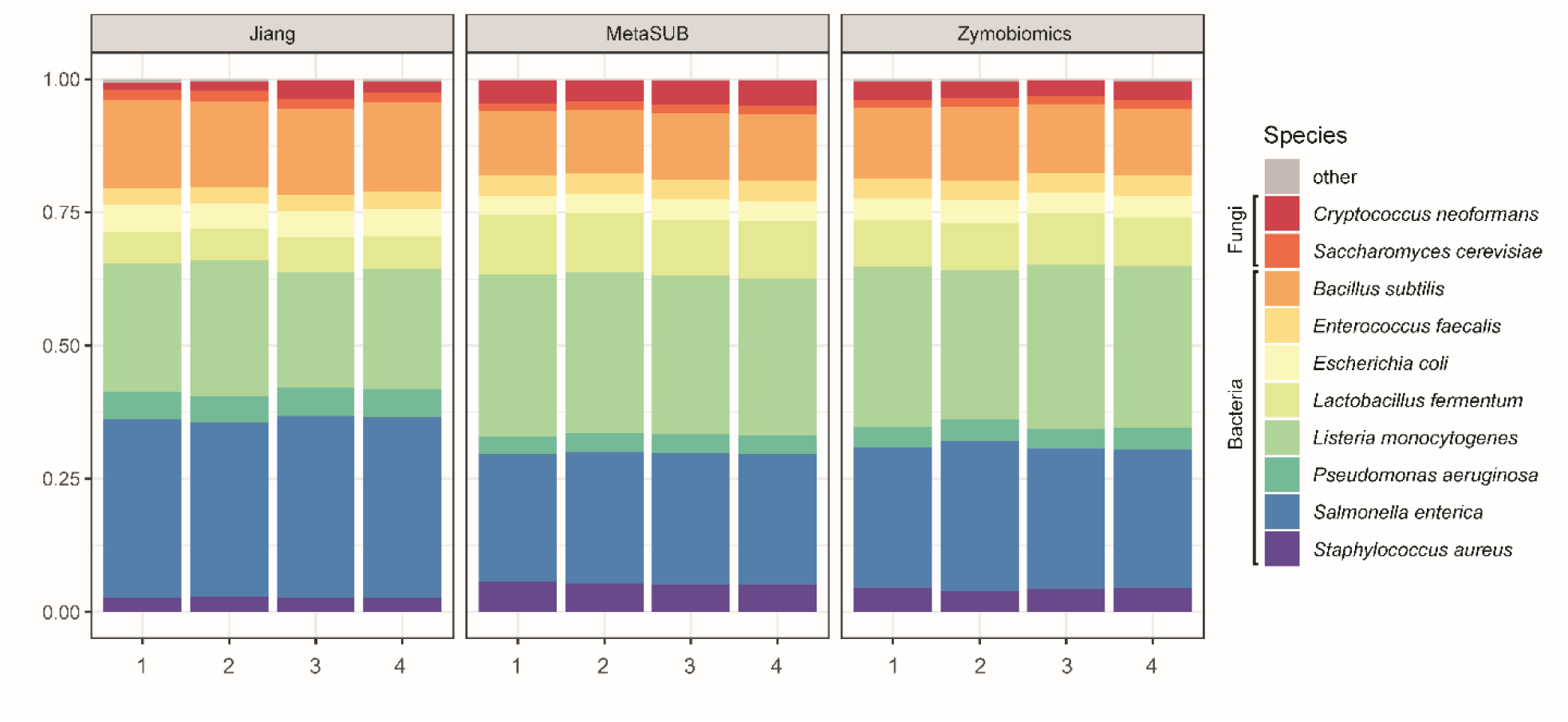
Relative distribution of the ten mock microbial community species for MetaSUB, Jiang, and Zymobiomics.

**Table 2.** Benchmarking results for MetaSUB, Jiang, and Zymobiomics on mock microbial community and subway air samples.

| A) Mock microbial community |  |  |  |  |  |  |
| --- | --- | --- | --- | --- | --- | --- |
| Measure | Within-sample differences | Est | 95% CI | T | df | P |
| Total DNA yield (ng) | MetaSUB – Jiang | -3 | [-28.5 , 22.5] | -0.37 | 3 | 0.73 |
|  | MetaSUB – Zymobiomics | -7 | [-46.5, 32.5] | -0.45 | 5 | 0.67 |
| 16S rRNA gene copy yield (copies) | MetaSUB – Jiang | 17107 | [-634, 34848] | 3.07 | 3 | 0.055 |
|  | MetaSUB – Zymobiomics | -11452 | [-83155, 60251] | -0.41 | 5 | 0.70 |

| B) Subway air samples |  |  |  |  |  |  |
| --- | --- | --- | --- | --- | --- | --- |
| Measure | Within-sample differences | Est | 95% CI | T | df | P |
| Total DNA yield (ng) | MetaSUB – Jiang | 1.07 | [0.77, 1.37] | 8.01 | 9 | < 0.001 |
|  | MetaSUB – Zymobiomics | 1.35 | [0.86, 1.85] | 5.94 | 13 | < 0.001 |
| 16S rRNA gene copy yield (copies) | MetaSUB – Jiang | 5046 | [3882, 6211] | 9.80 | 9 | < 0.001 |
|  | MetaSUB – Zymobiomics | 3451 | [1741, 5162] | 4.36 | 13 | < 0.001 |

One-sample t-test on within-sample differences ( $H_0$ : difference in within-sample measurements = 0) for different method pairs with mock microbial community (A) and subway air samples (B). Measures from Jiang/Zymobiomics were subtracted from the MetaSUB measures: the estimate (*est*) gives the departure from zero of the resultant values (larger than zero values indicate that MetaSUB had a higher yield than Jiang/Zymobiomics).

### Performance evaluation using subway air samples

The total DNA and 16S rRNA gene copy yields from subway air samples showed that the MetaSUB method obtained significantly higher total DNA and 16S rRNA gene copy yields than both Jiang and Zymobiomics (all *P* < 0.001; Figure 4; Table 2B).

The subway air samples that had been isolated with the MetaSUB method resulted in higher numbers of assigned reads than both Jiang (5 017 442 v. 2 630 115) and Zymobiomics (5 085 947 v. 4 601 016). Note that these results are average numbers from six individual air samples that were split and processed with the different method pairs, MetaSUB (N=3) v. Jiang (N=3) and MetaSUB (N=3) v. Zymobiomics (N=3). All samples reached saturation with regard to α-diversity at the lowest common assigned read depth (1 160 976, Figure S1), which was the depth at which all samples were rarified to. Taxonomic distributions at the family level were highly similar between the samples processed with MetaSUB and Zymobiomics (Figure 5). The samples processed with MetaSUB and Jiang were also highly similar, but a skew was observed in the relative abundances for two of the three Jiang samples (Figure 5). In the MetaSUB v. Zymobiomics comparison, the top ten most abundant phyla were identical between the method pairs, but not identical in their ordering by abundance (Table 3). Of the top ten families, one was uniquely found in the MetaSUB results (*Staphylococcaceae*; lowest abundance) and one in the Zymobiomics results (*Rhodobacteraceae*; second lowest abundance; Table 3). Among the ten top genera, only two where unique for MetaSUB (*Hymenobacter* and *Staphylococcus*) and two for Zymobiomics (*Dietzia* and *Paracoccus*; Table 3). Among the top ten species in each group, only one was unique to MetaSUB (*Chlorogloea sp. CCALA 695*) and one to Zymobiomics (*Lecanicillium sp. LEC01*; Table 3). In the MetaSUB v. Jiang comparison, there were more pronounced differences. The top ten phyla were not identical; *Acidobacteria* was only found in the MetaSUB results and *Planctomycetes* only in the Jiang results (Table 4). The top ten families were identical (but not in ordering); however, Jiang reported a substantially higher relative abundance of the family that was most abundant for both methods (*Micrococcaceae*, MetaSUB: 14% and Jiang: 25.6%; Table 4). Among the ten top genera, two where unique for MetaSUB (*Corynebacterium* and *Hymenobacter*) and two for Jiang (*Dietzia* and *Marmoricola*; Table 4). Here, the most abundant genus in Jiang (*Micrococcus*: 11.7%) was not the most abundant in MetaSUB (second most abundant; 5.60%) Among the top ten species in each group, only five species were present in both MetaSUB and Jiang results (Table 4).

**Figure 4.**
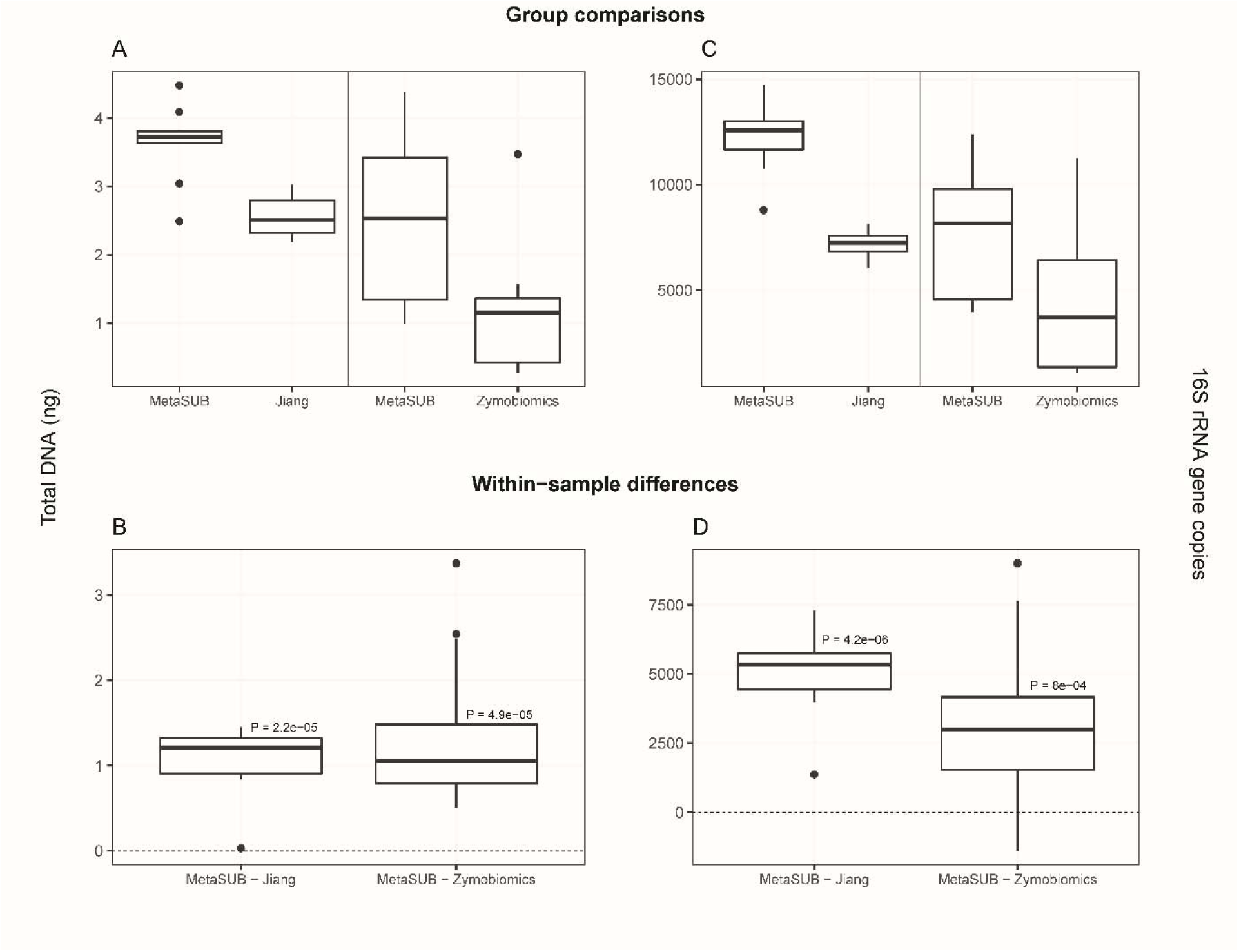
Benchmarking results for MetaSUB, Jiang, and Zymobiomics on split subway air samples. One sample t-tests were performed on within-sample differences (B, D) of total DNA yield (A), and 16S rRNA gene copy yield (C) for MetaSUB (N=10) and Jiang (N=10), and MetaSUB (N=14) and Zymobiomics (N=14).

**Figure 5.**
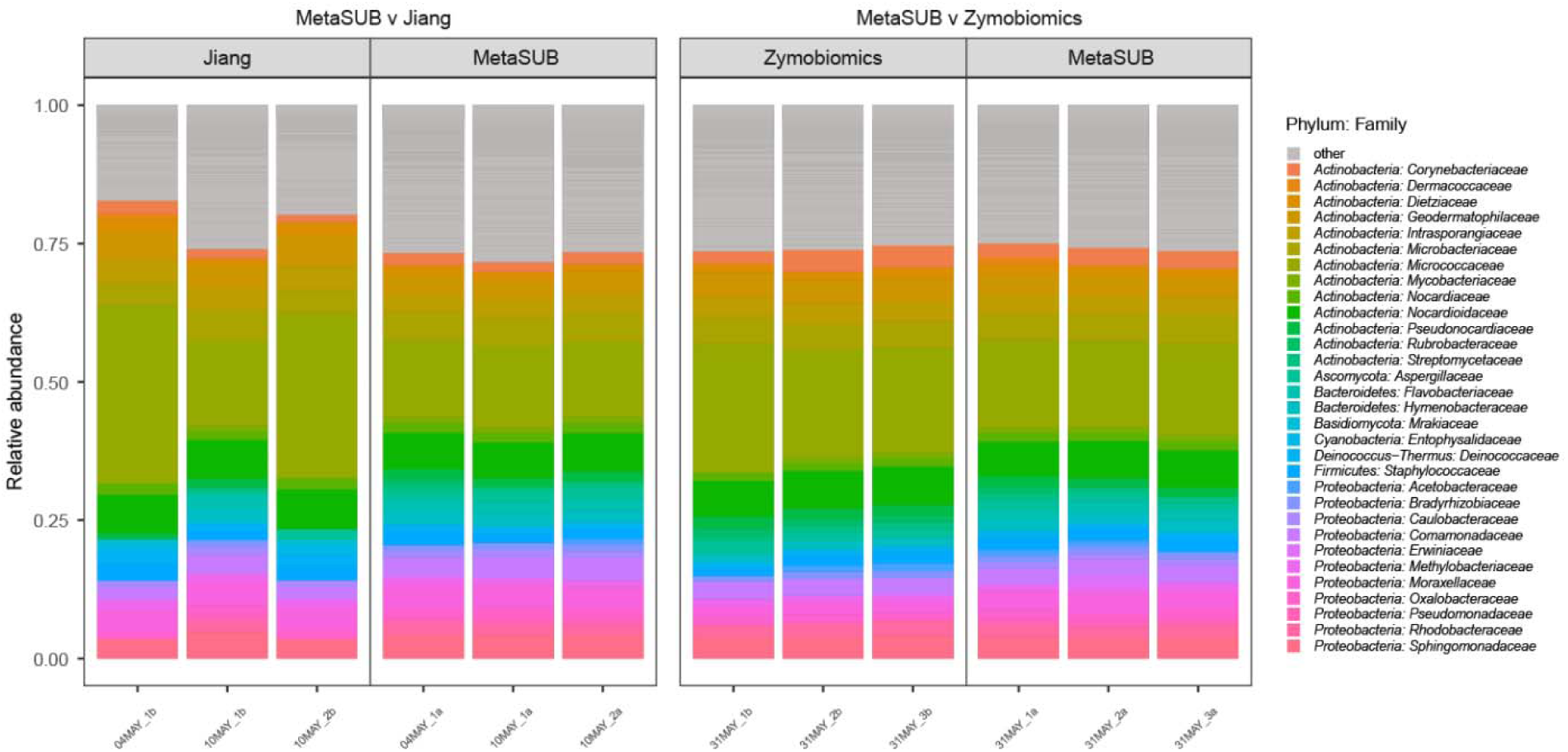
Relative taxonomic (family-level) distribution in split subway air samples (MetaSUB v. Jiang, MetaSUB v. Zymobiomics). Relative taxonomic (family-level) distribution in subway air samples (N=6) that were split and processed with the MetaSUB (N=3) and Jiang (N=3) or MetaSUB (N=3) and Zymobiomics (N=3) methods. Families with <1% representation are listed as “other”.

**Table 3.** Abundant microbial taxa in subway air samples (MetaSUB v. Zymobiomics method).

| MetaSUB v. Zymobiomics |  |  |  |  |  |  |  |
| --- | --- | --- | --- | --- | --- | --- | --- |
| MetaSub |  |  |  | Zymobiomics |  |  |  |
| Phylum | prevalence<br>mean | total | abundance | Phylum | prevalence<br>mean | total | Abundance |
| *Actinobacteria* | 2.7 | 9271 | 51.61 % | *Actinobacteria* | 2.7 | 9301 | 57.72 % |
| *Proteobacteria* | 2.3 | 14269 | 27.02 % | *Proteobacteria* | 2.2 | 13421 | 23.67 % |
| *Firmicutes* | 1.6 | 3250 | 4.76 % | *Ascomycota* | 2.5 | 1899 | 4.91 % |
| *Bacteroidetes* | 2.0 | 2999 | 4.74 % | *Basidiomycota* | 2.4 | 622 | 4.75 % |
| *Ascomycota* | 2.4 | 1843 | 4.42 % | *Firmicutes* | 1.3 | 2705 | 3.30 % |
| *Basidiomycota* | 2.3 | 606 | 3.84 % | *Bacteroidetes* | 1.5 | 2257 | 2.77 % |
| *Cyanobacteria* | 2.3 | 526 | 1.35 % | *Deinococcus-Thermus* | 2.6 | 204 | 1.26 % |
| *Deinococcus-Thermus* | 2.5 | 196 | 1.23 % | *Cyanobacteria* | 1.9 | 432 | 0.66 % |
| *Euryarchaeota* | 1.7 | 450 | 0.62 % | *Euryarchaeota* | 1.5 | 393 | 0.58 % |
| *Acidobacteria* | 2.5 | 112 | 0.08 % | *Acidobacteria* | 2.6 | 114 | 0.08 % |
| Family | prevalence<br>mean | total | abundance | Family | prevalence<br>mean | total | Abundance |
| *Micrococcaceae* | 2.8 | 801 | 15.72 % | *Micrococcaceae* | 2.8 | 822 | 20.34 % |
| *Nocardiodaceae* | 2.8 | 308 | 6.51 % | *Nocardiodaceae* | 2.9 | 311 | 6.65 % |
| *Microbacteriaceae* | 2.6 | 1222 | 4.89 % | *Microbacteriaceae* | 2.6 | 1235 | 4.70 % |
| *Sphingomonadaceae* | 2.8 | 1006 | 3.88 % | *Geodermatophilaceae* | 3.0 | 150 | 4.24 % |
| *Geodermatophilaceae* | 3.0 | 150 | 3.67 % | *Sphingomonadaceae* | 2.8 | 1013 | 4.04 % |
| *Moraxellaceae* | 1.9 | 608 | 3.54 % | *Intrasporangiaceae* | 3.0 | 211 | 3.46 % |
| *Intrasporangiaceae* | 3.0 | 211 | 3.15 % | *Corynebacteriaceae* | 2.6 | 571 | 3.35 % |
| *Comamonadaceae* | 2.6 | 800 | 2.94 % | *Comamonadaceae* | 2.7 | 838 | 3.02 % |
| *Corynebacteriaceae* | 2.6 | 568 | 2.93 % | *Rhodobacteraceae* | 2.6 | 1484 | 2.28 % |
| *Staphylococcaceae* | 2.3 | 401 | 2.17 % | *Moraxellaceae* | 1.6 | 533 | 2.14 % |
| Genus | prevalence<br>mean | total | abundance | Genus | prevalence<br>mean | total | Abundance |
| *Micrococcus* | 2.9 | 29 | 7.27 % | *Micrococcus* | 2.9 | 29 | 9.34 % |
| *Arthrobacter* | 2.8 | 399 | 5.32 % | *Arthrobacter* | 2.9 | 413 | 6.44 % |
| *Nocardioide* | 2.8 | 176 | 3.51 % | *Nocardioide* | 2.9 | 177 | 3.57 % |
| *Sphingomonas* | 2.9 | 461 | 3.09 % | *Kocuria* | 2.7 | 79 | 3.44 % |
| *Corynebacterium* | 2.6 | 568 | 2.93 % | *Corynebacterium* | 2.6 | 571 | 3.35 % |
| *Psychrobacter* | 2.0 | 141 | 2.85 % | *Sphingomonas* | 2.9 | 464 | 3.23 % |
| *Blastococcus* | 3.0 | 60 | 2.44 % | *Blastococcus* | 3.0 | 60 | 2.92 % |
| *Staphylococcus* | 2.2 | 308 | 2.02 % | *Psychrobacter* | 2.7 | 184 | 1.82 % |
| *Kocuria* | 2.7 | 79 | 2.02 % | *Dietzia* | 3.0 | 51 | 1.70 % |
| *Hymenobacter* | 2.9 | 98 | 1.92 % | *Paracoccus* | 2.9 | 167 | 1.58 % |
| Species | prevalence<br>mean | total | abundance | Species | prevalence<br>mean | total | Abundance |
| *Micrococcus luteus* | 3.0 | 3 | 1.13 % | *Micrococcus luteus* | 3.0 | 3 | 1.44 % |
| *Arthrobacter sp. H41* | 3.0 | 3 | 1.00 % | *Arthrobacter sp. H41* | 3.0 | 3 | 1.38 % |
| *Rubrobacter aphysinae* | 3.0 | 3 | 0.78 % | *Rubrobacter aphysinae* | 3.0 | 3 | 0.97 % |
| *Arthrobacter sp. Leaf234* | 3.0 | 3 | 0.67 % | *Arthrobacter sp. Leaf234* | 3.0 | 3 | 0.89 % |
| *Marmoricola sp. Leaf446* | 3.0 | 3 | 0.65 % | *Stereum hirsutum* | 3.0 | 3 | 0.76 % |
| *Chlorogloea sp. CCA 695* | 3.0 | 3 | 0.63 % | *Marmoricola sp. Leaf446* | 3.0 | 3 | 0.76 % |
| *Deinococcus marmoris* | 3.0 | 3 | 0.60 % | *Fomitopsis pinicola* | 3.0 | 3 | 0.68 % |
| *Stereum hirsutum* | 3.0 | 3 | 0.50 % | *Blastococcus sp. DSM 44268* | 3.0 | 3 | 0.64 % |
| *Blastococcus sp. DSM 44268* | 3.0 | 3 | 0.48 % | *Deinococcus marmoris* | 3.0 | 3 | 0.57 % |
| *Fomitopsis pinicola* | 3.0 | 3 | 0.45 % | *Lecanicillium sp. LEC01* | 3.0 | 3 | 0.57 % |

Top ten microbial phyla, families, genera and species in subway air samples (N=3) that were split
and processed with the MetaSUB (N=3) and Zymobiomics (N=3) methods.

**Table 4.** Abundant microbial taxa in subway air samples (MetaSUB v. Jiang method).

| MetaSUB v. Jiang |  |  |  |  |  |  |  |
| --- | --- | --- | --- | --- | --- | --- | --- |
| MetaSub |  |  |  | Jiang |  |  |  |
| Phylum | prevalence<br>mean | total | abundance | Phylum | prevalence<br>mean | total | Abundance |
| *Actinobacteria* | 2.7 | 9248 | 48.69 % | *Actinobacteria* | 1.1 | 3625 | 60.08 % |
| *Proteobacteria* | 2.3 | 14379 | 28.58 % | *Proteobacteria* | 1.0 | 6090 | 23.40 % |
| *Ascomycota* | 2.4 | 1843 | 5.28 % | *Firmicutes* | 0.7 | 1510 | 4.58 % |
| *Bacteroidetes* | 2.1 | 3148 | 5.10 % | *Bacteroidetes* | 0.9 | 1436 | 3.40 % |
| *Firmicutes* | 1.6 | 3250 | 4.52 % | *Ascomycota* | 1.1 | 822 | 2.78 % |
| *Basidiomycota* | 2.3 | 598 | 3.95 % | *Deinococcus-Thermus* | 1.0 | 77 | 1.94 % |
| *Cyanobacteria* | 2.3 | 518 | 1.44 % | *Basidiomycota* | 1.1 | 286 | 1.91 % |
| *Deinococcus-Thermus* | 2.4 | 190 | 1.35 % | *Cyanobacteria* | 1.1 | 243 | 1.50 % |
| *Euryarchaeota* | 1.9 | 495 | 0.63 % | *Euryarchaeota* | 0.9 | 231 | 0.15 % |
| *Acidobacteria* | 2.6 | 113 | 0.08 % | *Planctomycetes* | 1.2 | 52 | 0.04 % |
| Family | prevalence<br>mean | total | abundance | Family | prevalence<br>mean | total | abundance |
| *Micrococcaceae* | 2.7 | 785 | 13.96 % | *Micrococcaceae* | 1.2 | 336 | 25.63 % |
| *Nocardiodaceae* | 2.8 | 301 | 6.58 % | *Nocardiodaceae* | 1.1 | 123 | 7.02 % |
| *Microbacteriaceae* | 2.6 | 1213 | 5.05 % | *Geodermatophilaceae* | 1.3 | 65 | 4.77 % |
| *Sphingomonadaceae* | 2.8 | 1010 | 4.20 % | *Microbacteriaceae* | 1.0 | 486 | 4.45 % |
| *Moraxellaceae* | 2.0 | 655 | 4.12 % | *Intrasporangiaceae* | 1.4 | 97 | 4.15 % |
| *Comamonadaceae* | 2.4 | 750 | 3.68 % | *Moraxellaceae* | 0.9 | 289 | 4.05 % |
| *Geodermatophilaceae* | 3.0 | 149 | 3.59 % | *Sphingomonadaceae* | 1.0 | 357 | 3.92 % |
| *Intrasporangiaceae* | 3.0 | 210 | 3.23 % | *Comamonadaceae* | 1.1 | 337 | 2.60 % |
| *Hymenobacteraceae* | 2.9 | 182 | 2.17 % | *Staphylococcaceae* | 0.9 | 169 | 2.39 % |
| *Flavobacteriaceae* | 2.2 | 1594 | 2.04 % | *Dietziaceae* | 1.1 | 19 | 1.99 % |
| Genus | prevalence<br>mean | total | abundance | Genus | prevalence<br>mean | total | abundance |
| *Arthrobacter* | 2.7 | 391 | 5.63 % | *Micrococcus* | 2.1 | 21 | 11.69 % |
| *Micrococcus* | 2.8 | 28 | 5.60 % | *Arthrobacter* | 1.1 | 153 | 9.34 % |
| *Nocardioide* | 2.7 | 167 | 3.50 % | *Kocuria* | 1.2 | 34 | 3.53 % |
| *Psychrobacter* | 1.9 | 131 | 3.40 % | *Psychrobacter* | 0.9 | 64 | 3.46 % |
| *Sphingomonas* | 2.9 | 465 | 3.38 % | *Nocardioide* | 1.0 | 61 | 3.38 % |
| *Blastococcus* | 3.0 | 59 | 2.30 % | *Sphingomonas* | 1.0 | 155 | 3.20 % |
| *Corynebacterium* | 2.6 | 559 | 2.04 % | *Blastococcus* | 1.3 | 26 | 3.16 % |
| *Hymenobacter* | 2.9 | 98 | 2.02 % | *Marmoricola* | 2.0 | 12 | 2.78 % |
| *Staphylococcus* | 2.0 | 289 | 1.77 % | *Staphylococcus* | 0.9 | 128 | 2.33 % |
| *Kocuria* | 2.7 | 79 | 1.76 % | *Dietzia* | 1.1 | 19 | 1.99 % |
| Species | prevalence<br>mean | total | abundance | Species | prevalence<br>mean | total | abundance |
| *Arthrobacter* sp. H41 | 3.0 | 3 | 1.14 % | *Arthrobacter* sp. H41 | 3.0 | 3 | 2.36 % |
| *Micrococcus luteus* | 3.0 | 3 | 0.88 % | *Arthrobacter* sp. Leaf234 | 3.0 | 3 | 2.15 % |
| *Chlorogloea* sp. CCALA 695 | 3.0 | 3 | 0.74 % | *Marmoricola* sp. Leaf446 | 3.0 | 3 | 1.71 % |
| *Rubrobacter aplysinae* | 3.0 | 3 | 0.71 % | *Deinococcus marmoris* | 3.0 | 3 | 1.56 % |
| *Arthrobacter* sp. Leaf234 | 3.0 | 3 | 0.71 % | *Blastococcus* sp. DSM 44268 | 3.0 | 3 | 1.24 % |
| *Aspergillus* sp. MA 6041 | 3.0 | 3 | 0.69 % | *Arthrobacter agilis* | 3.0 | 3 | 1.23 % |
| *Marmoricola* sp. Leaf446 | 3.0 | 3 | 0.67 % | *Chlorogloea* sp. CCALA 695 | 3.0 | 3 | 1.14 % |
| *Deinococcus marmoris* | 3.0 | 3 | 0.66 % | *Marmoricola scoriae* | 3.0 | 3 | 0.84 % |
| *Acidovorax temperans* | 3.0 | 3 | 0.61 % | *Janibacter* sp. Soil728 | 3.0 | 3 | 0.82 % |
| *Stereum hirsutum* | 3.0 | 3 | 0.57 % | *Mrakia frigida* | 3.0 | 3 | 0.77 % |

Top ten microbial phyla, families, genera and species in subway air samples (N=3) that were split
and processed with the MetaSUB (N=3) and Jiang (N=3) methods.

Linear regression of within-sample α-diversity indices showed that MetaSUB reported significantly higher diversity estimates compared to Zymobiomics (Observed: *est*=734.3, *P*=0.01; Shannon: *est*=0.22, *P*=0.002; Simpson: *est*=0.00079, *P*=0.001; Figure 6), but no differences were shown between MetaSUB and Jiang α-diversity estimates (Observed: *est*=6531; Shannon: *est*=2.75; Simpson: *est*=0.028; all *P*>0.12; Figure 6). PERMANOVA tests of PCoA ordinated Bray Curtis distances found no significant differences among MetaSUB and Jiang (*P*=0.1) or MetaSUB and Zymobiomics (*P*=0.1; Figure 7).

**Figure 6.**
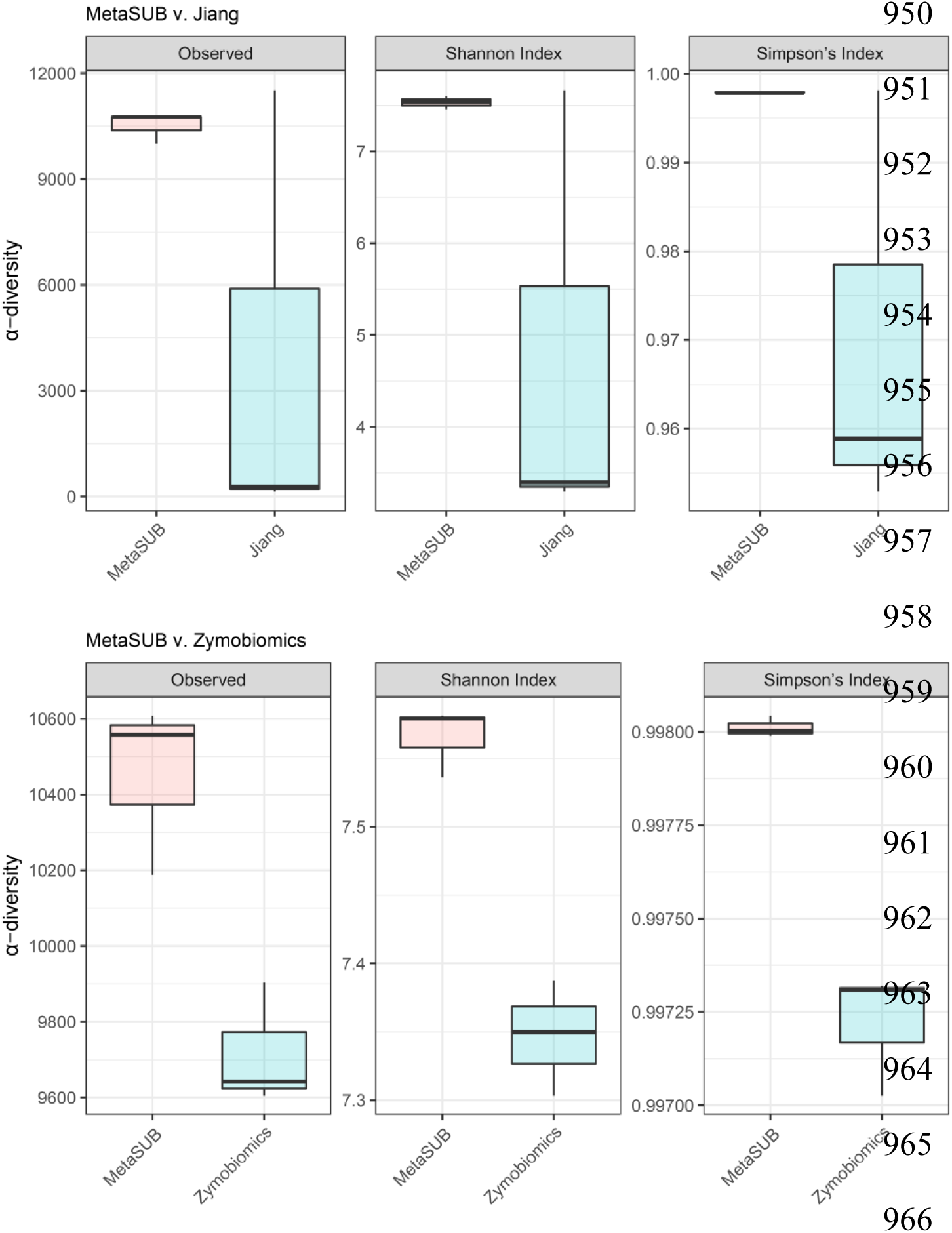
Diversity estimates (α-diversity) for split subway air samples (MetaSUB v. Jiang, MetaSUB v. Zymobiomics). Comparison of diversity estimates (α-diversity) for subway air samples (N=6) that were split and processed with the MetaSUB (N=3) and Jiang (N=3) or MetaSUB (N=3) and Zymobiomics (N=3) methods.

**Figure 7.**
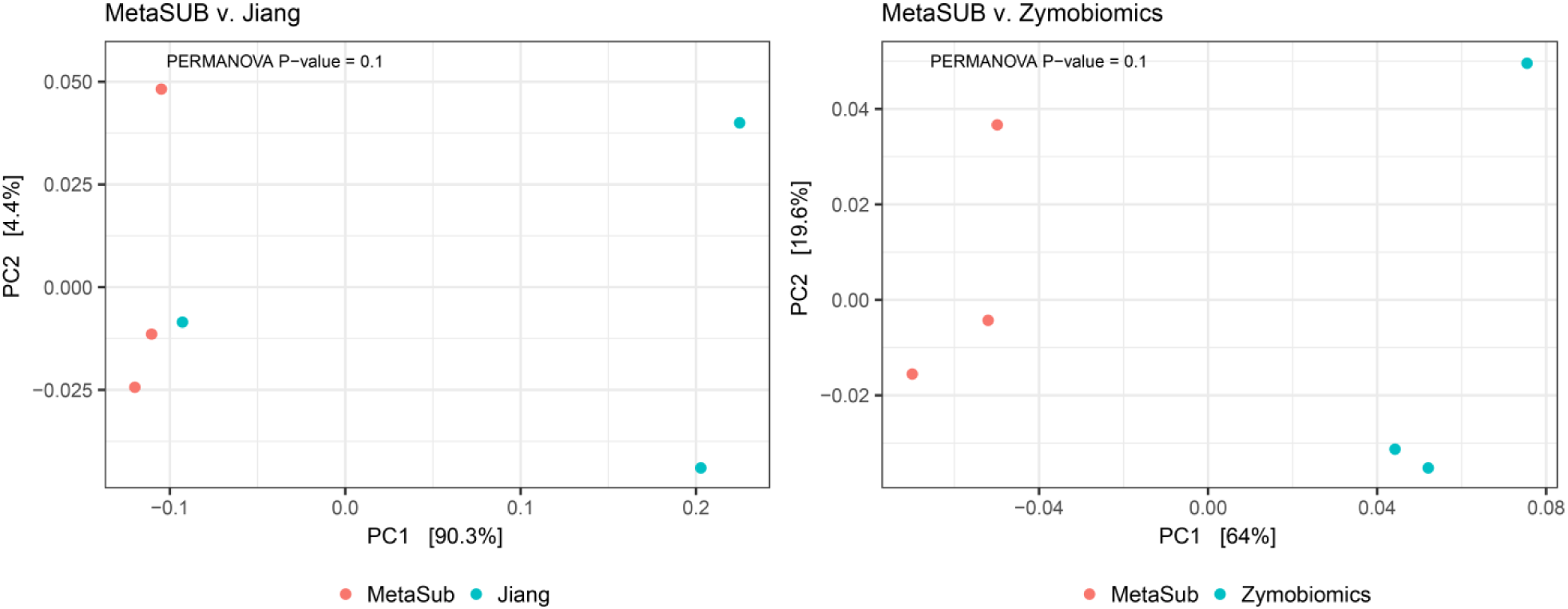
PCoA ordination plots (β-diversity) for split subway air samples (MetaSUB v. Jiang, MetaSUB v. Zymobiomics). PCoA ordination plots using Bray Curtis distance estimation (β-diversity) for subway air samples (N=6) that were split and processed with the MetaSUB (N=3) and Jiang (N=3) or MetaSUB (N=3) and Zymobiomics (N=3) methods. PERMANOVA tests were performed on the MetaSUB/Jiang and MetaSUB/Zymobiomics groupings.

The random forest classification analysis, where species-level features were scored by their ability to correctly classify the DNA isolation method used, had a perfect out-of-bag error of 0%, and a significant permutation test (*P*>0.02) for MetaSUB v. Zymobiomics. For MetaSUB v. Jiang, the classification model had an out-of-bag error of 16%, but also here the permutation test was significant (*P*=0.01). For MetaSUB v. Zymobiomics, the proportions of archaea, bacteria and fungi across the dataset and in the 100 species most important for correctly classifying samples as either MetaSUB or Zymobiomics were highly similar. However, for MetaSUB v. Jiang, 6.0% of all assigned species were fungi, while among the 100 species most important for classification, 20 were fungi. These 20 fungal species all had higher abundances in the MetaSUB results (Figure S4). The top 30 most important features for both classification models are shown in Figure 8.

**Figure 8.**
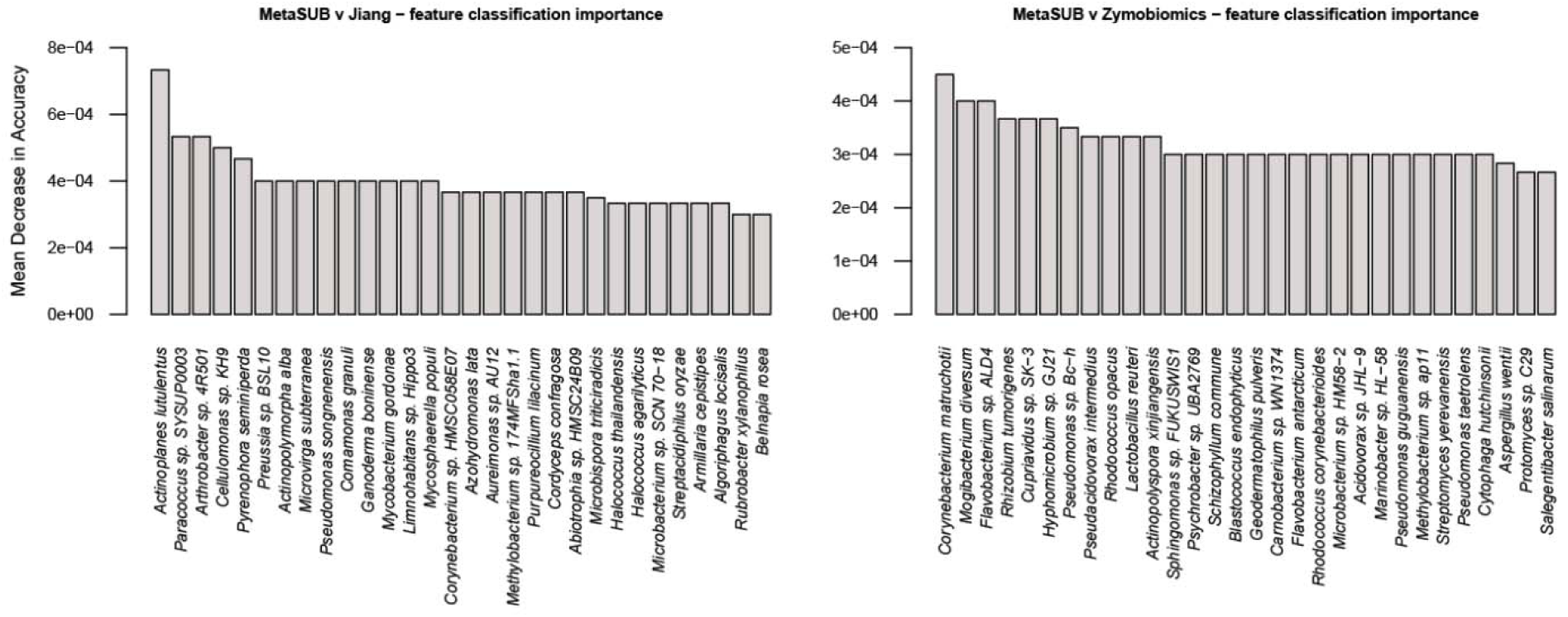
Random forest classification analysis of split subway air samples (MetaSUB v. Jiang, MetaSUB v. Zymobiomics). Random forest classification analysis of subway air samples (N=6) that were split and processed with the MetaSUB (N=3) and Jiang (N=3) or MetaSUB (N=3) and Zymobiomics (N=3) methods, showing taxonomic features with the highest classification variable importance for correctly identifying the DNA isolation method.

### DNA distribution in intermediate pellet/supernatant fractions of the MetaSUB method

The distribution of DNA in terms of both amount and composition (diversity) in the intermediate pellet and supernatant fractions of the MetaSUB method was investigated by separately isolating DNA from the two fractions from subway air samples. The results revealed that the supernatant fraction contained 42%±6 of the total DNA yield and 32%±12 of the total 16S rRNA gene copy yield (Figure S2).

The SMS results showed that the pellet samples had a higher number of assigned reads than supernatant samples (2 584 159 v. 1 609 457). Rarefication plots of pellet and supernatant samples indicated that α-diversity indices (particularly Shannon and Simpson) reached saturation before the lowest common assigned read depth (453 218, Figure S2), which was the depth at which all samples were rarified to. The taxonomic distributions in pellet and supernatant samples were largely similar (Table 5; Figure 9). The top ten phyla were identical in the pellet and supernatant group, but not identical in their ordering by abundance (Table 5). Of the top ten families, one was uniquely found in the pellet group (*Rhodobacteraceae*; second lowest abundance) and one only in the supernatant group (*Deinococcaceae*; lowest abundance; Table 5). Among the ten top genera, only one was unique for the pellet group (*Marmoricola*) and one for the supernatant group (*Deinococcus*; Table 5). Among the top ten species in each group, seven species were present in both (Table 5). Linear regression of within-sample α-diversity indices revealed no significant differences between pellet and supernatant samples (Figure 10; all *P*>0.38). A PERMANOVA test of PCoA ordinated Bray Curtis distances found that whether samples were pellet or supernatant explained 51.7% of the among-sample variance in diversity (Figure 11; *P*=0.004).

**Figure 9.**
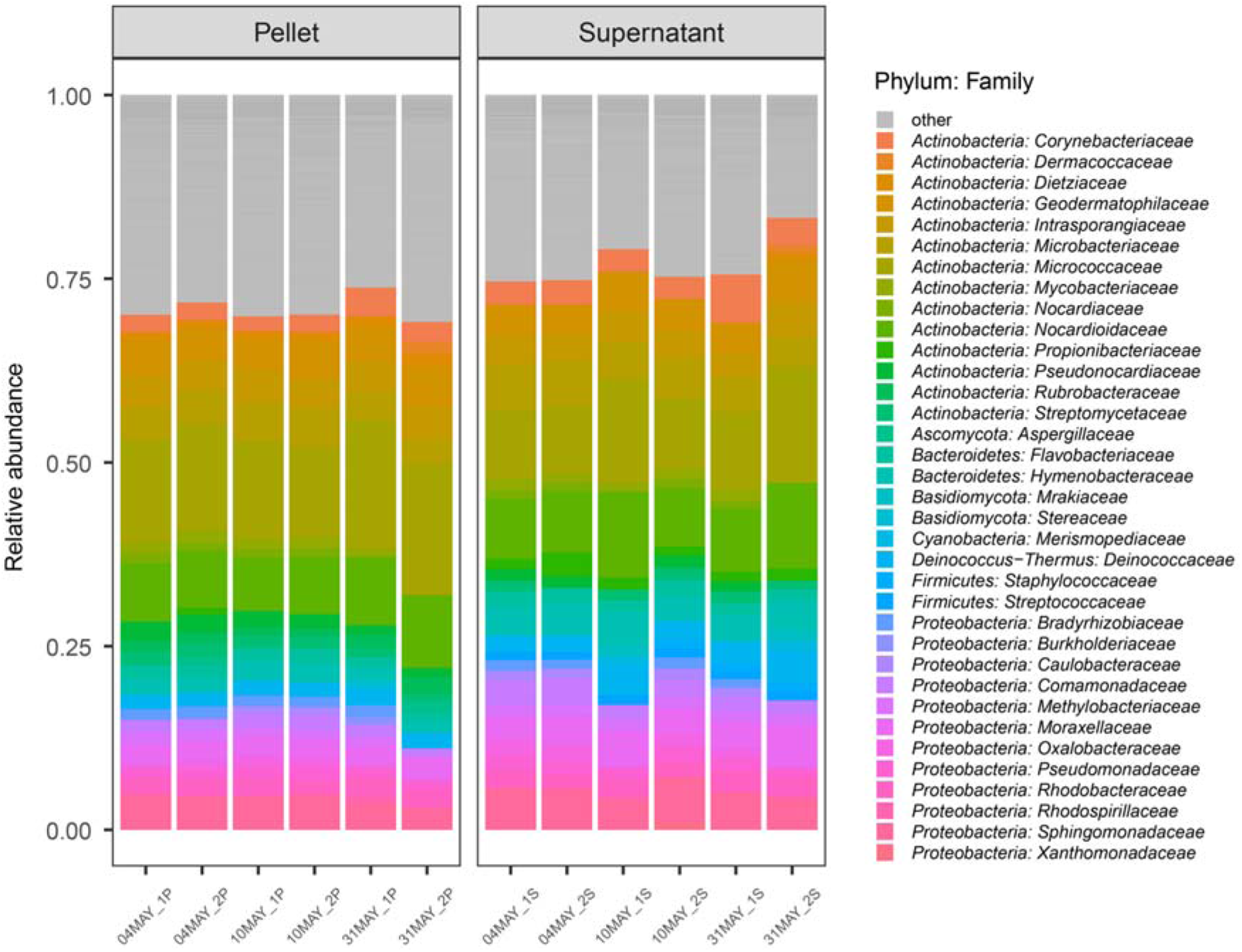
Relative taxonomic (family-level) distribution in pellet and supernatant fractions from subway air samples (MetaSUB method). Relative taxonomic (family-level) distribution for subway air samples (N=6) where the intermediate pellet (N=6) and supernatant (N=6) fractions were processed separately with the MetaSUB method. Families with <1% representation are listed as “other”.

**Figure 10.**
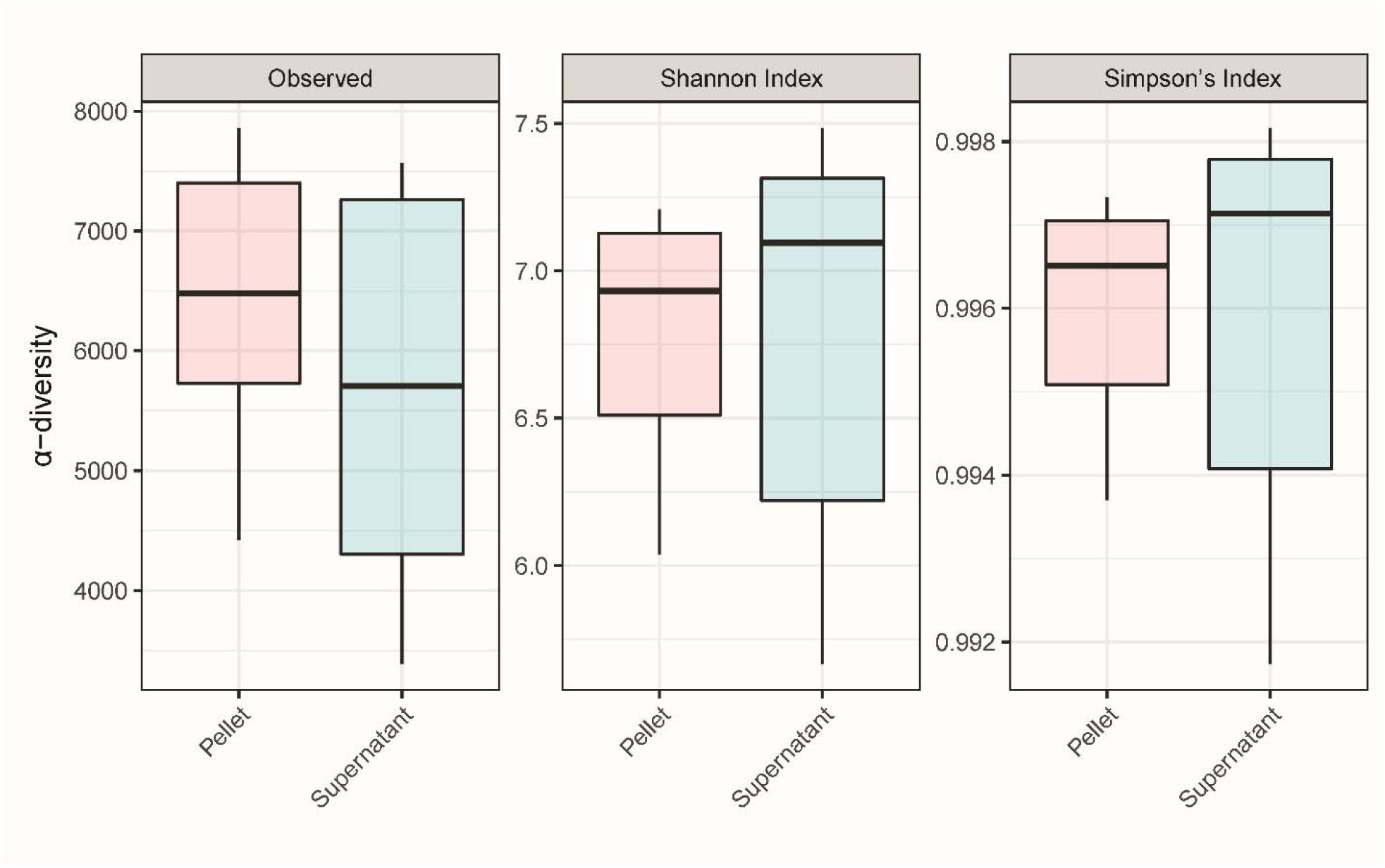
Diversity estimates (α-diversity) for pellet and supernatant fractions from subway air samples (MetaSUB method). Diversity estimates (α-diversity) for subway air samples (N=6) where the intermediate pellet (N=6) and supernatant (N=6) fractions were processed separately with the MetaSUB method.

**Figure 11.**
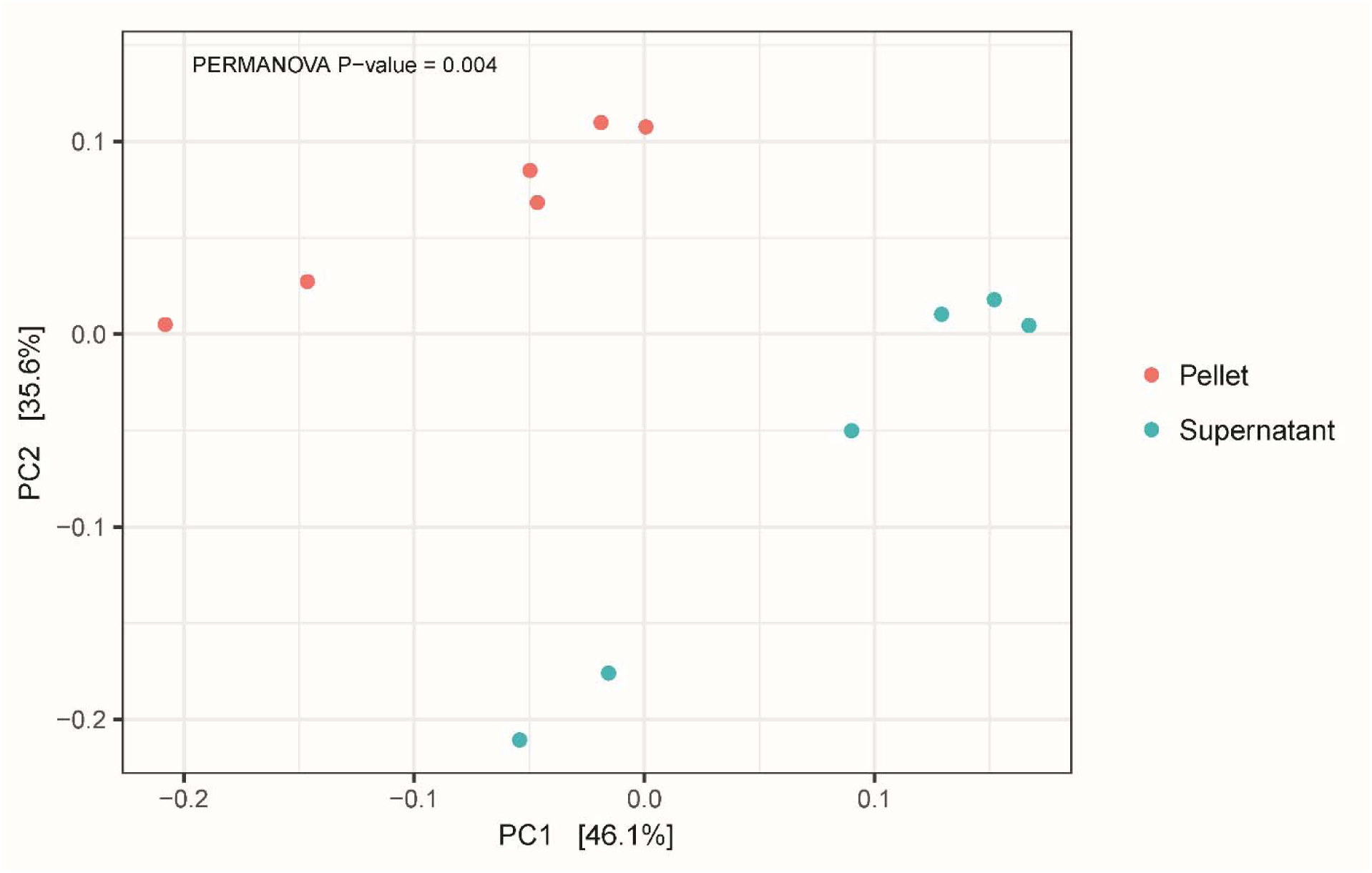
PCoA ordination plot (β-diversity) for pellet and supernatant fractions from subway air samples (MetaSUB method). PCoA ordination plot using Bray Curtis distance estimation (β-diversity) for subway air samples (N=6) where the intermediate pellet (N=6) and supernatant (N=6) fractions were processed separately with the MetaSUB method. PERMANOVA test was performed on pellet/supernatant grouping.

**Table 5.** Abundant microbial taxa in pellet and supernatant fractions from subway air samples (MetaSUB method).

| Supernatant |  |  |  | Pellet |  |  |  |
| --- | --- | --- | --- | --- | --- | --- | --- |
| Phylum | Prevalence<br>mean | Total | abundance | Phylum | Prevalence<br>mean | Total | abundance |
| *Actinobacteria* | 3.5 | 9479 | 51.21% | *Actinobacteria* | 4.0 | 10653 | 53.53% |
| *Proteobacteria* | 3.1 | 14453 | 28.87% | *Proteobacteria* | 3.4 | 15894 | 24.50% |
| *Bacteroidetes* | 2.8 | 3139 | 7.10% | *Basidiomycota* | 3.7 | 717 | 5.42% |
| *Firmicutes* | 2.5 | 3816 | 4.23% | *Ascomycota* | 4.2 | 2109 | 5.23% |
| *Deinococcus-Thermus* | 3.2 | 184 | 3.06% | *Bacteroidetes* | 2.9 | 3202 | 4.44% |
| *Basidiomycota* | 1.5 | 290 | 1.86% | *Firmicutes* | 2.6 | 3874 | 2.68% |
| *Ascomycota* | 2.4 | 1231 | 1.62% | *Deinococcus-Thermus* | 3.7 | 214 | 2.05% |
| *Cyanobacteria* | 2.8 | 431 | 1.47% | *Cyanobacteria* | 3.6 | 557 | 1.03% |
| *Euryarchaeota* | 1.5 | 260 | 0.17% | *Euryarchaeota* | 2.1 | 367 | 0.66% |
| *Acidobacteria* | 3.4 | 103 | 0.10% | *Acidobacteria* | 4.4 | 132 | 0.11% |
| Family | Prevalence<br>mean | Total | abundance | Family | Prevalence<br>mean | Total | abundance |
| *Micrococcaceae* | 3.6 | 735 | 11.27% | *Micrococcaceae* | 3.9 | 795 | 14.70% |
| *Nocardiodaceae* | 3.1 | 236 | 9.32% | *Nocardiodaceae* | 3.0 | 222 | 8.26% |
| *Microbacteriaceae* | 3.4 | 921 | 5.24% | *Geodermatophilaceae* | 2.9 | 97 | 4.65% |
| *Sphingomonadaceae* | 3.8 | 1119 | 5.20% | *Microbacteriaceae* | 3.5 | 958 | 4.53% |
| *Geodermatophilaceae* | 3.7 | 121 | 4.60% | *Sphingomonadaceae* | 4.3 | 1260 | 4.22% |
| *Moraxellaceae* | 2.9 | 794 | 4.14% | *Intrasporangiaceae* | 3.8 | 199 | 4.20% |
| *Hymenobacteraceae* | 3.5 | 150 | 3.93% | *Moraxellaceae* | 2.6 | 717 | 2.77% |
| *Intrasporangiaceae* | 3.7 | 192 | 3.81% | *Corynebacteriaceae* | 4.2 | 881 | 2.61% |
| *Corynebacteriaceae* | 3.7 | 789 | 3.77% | *Rhodobacteraceae* | 4.3 | 1698 | 2.33% |
| *Deinococcaceae* | 4.0 | 117 | 3.03% | *Hymenobacteraceae* | 3.8 | 163 | 2.15% |
| Genus | Prevalence<br>mean | Total | abundance | Genus | Prevalence<br>mean | Total | abundance |
| *Arthrobacter* | 3.3 | 320 | 6.31% | *Arthrobacter* | 3.7 | 357 | 7.74% |
| *Sphingomonas* | 3.7 | 512 | 4.40% | *Micrococcus* | 3.9 | 39 | 4.19% |
| *Nocardioides* | 3.0 | 127 | 3.87% | *Sphingomonas* | 4.1 | 560 | 3.40% |
| *Hymenobacter* | 3.4 | 89 | 3.85% | *Nocardioides* | 2.9 | 120 | 3.14% |
| *Corynebacterium* | 3.8 | 767 | 3.73% | *Blastococcus* | 3.5 | 39 | 3.13% |
| *Psychrobacter* | 2.5 | 143 | 3.24% | *Corynebacterium* | 4.2 | 854 | 2.59% |
| *Deinococcus* | 4.0 | 117 | 3.03% | *Marmoricola* | 5.8 | 23 | 2.56% |
| *Friedmanniella* | 6.0 | 18 | 2.85% | *Friedmanniella* | 6.0 | 18 | 2.38% |
| *Blastococcus* | 4.1 | 45 | 2.60% | *Psychrobacter* | 2.3 | 133 | 2.37% |
| *Micrococcus* | 4.6 | 46 | 2.31% | *Hymenobacter* | 3.2 | 84 | 2.04% |
| Species | Prevalence<br>mean | Total | abundance | Species | Prevalence<br>mean | Total | abundance |
| *Deinococcus marmoris* | 6.0 | 6 | 2.07% | *Arthrobacter sp. H41* | 6.0 | 6 | 2.07% |
| *Arthrobacter sp. Leaf234* | 6.0 | 6 | 1.91% | *Micrococcus luteus* | 6.0 | 6 | 1.96% |
| *Marmoricola sp. Leaf446* | 6.0 | 6 | 1.31% | *Rubrobacter aphysinae* | 6.0 | 6 | 1.52% |
| *Friedmanniella flava* | 6.0 | 6 | 1.05% | *Arthrobacter sp. Leaf234* | 6.0 | 6 | 1.51% |
| *Friedmanniella sagamiharensis* | 6.0 | 6 | 1.03% | *Marmoricola sp. Leaf446* | 6.0 | 6 | 1.42% |
| *Cutibacterium acnes* | 6.0 | 6 | 1.02% | *Blastococcus sp. DSM 44268* | 6.0 | 6 | 1.32% |
| *Mrakia frigida* | 6.0 | 6 | 1.02% | *Deinococcus marmoris* | 6.0 | 6 | 1.27% |
| *Blastococcus sp. DSM 44268* | 6.0 | 6 | 1.02% | *Arthrobacter agilis* | 6.0 | 6 | 0.95% |
| *Micrococcus luteus* | 6.0 | 6 | 1.01% | *Friedmanniella flava* | 6.0 | 6 | 0.89% |
| *Arthrobacter sp. H41* | 6.0 | 6 | 0.98% | *Stereum hirsutum* | 6.0 | 6 | 0.86% |

Top ten microbial phyla, families, genera and species in the intermediate pellet (N=6) and supernatant (N=6) fractions from subway air samples (N=6) processed with the MetaSUB method.

The cross-kingdom analysis revealed substantial differences in the relative representation of almost all examined groups (archaea, bacteria, fungi, plants, human, and other animals) between the pellet and supernatant samples (Figure 12). While very few reads were assigned to archaea, only pellet samples had any coverage within this group. Pellet samples also had a higher relative number of assigned reads across all sample pairs within bacteria and fungi. The supernatant had a higher relative number of reads assigned as human and other animals, while plants saw similar representation in pellet and supernatant samples.

**Figure 12.**
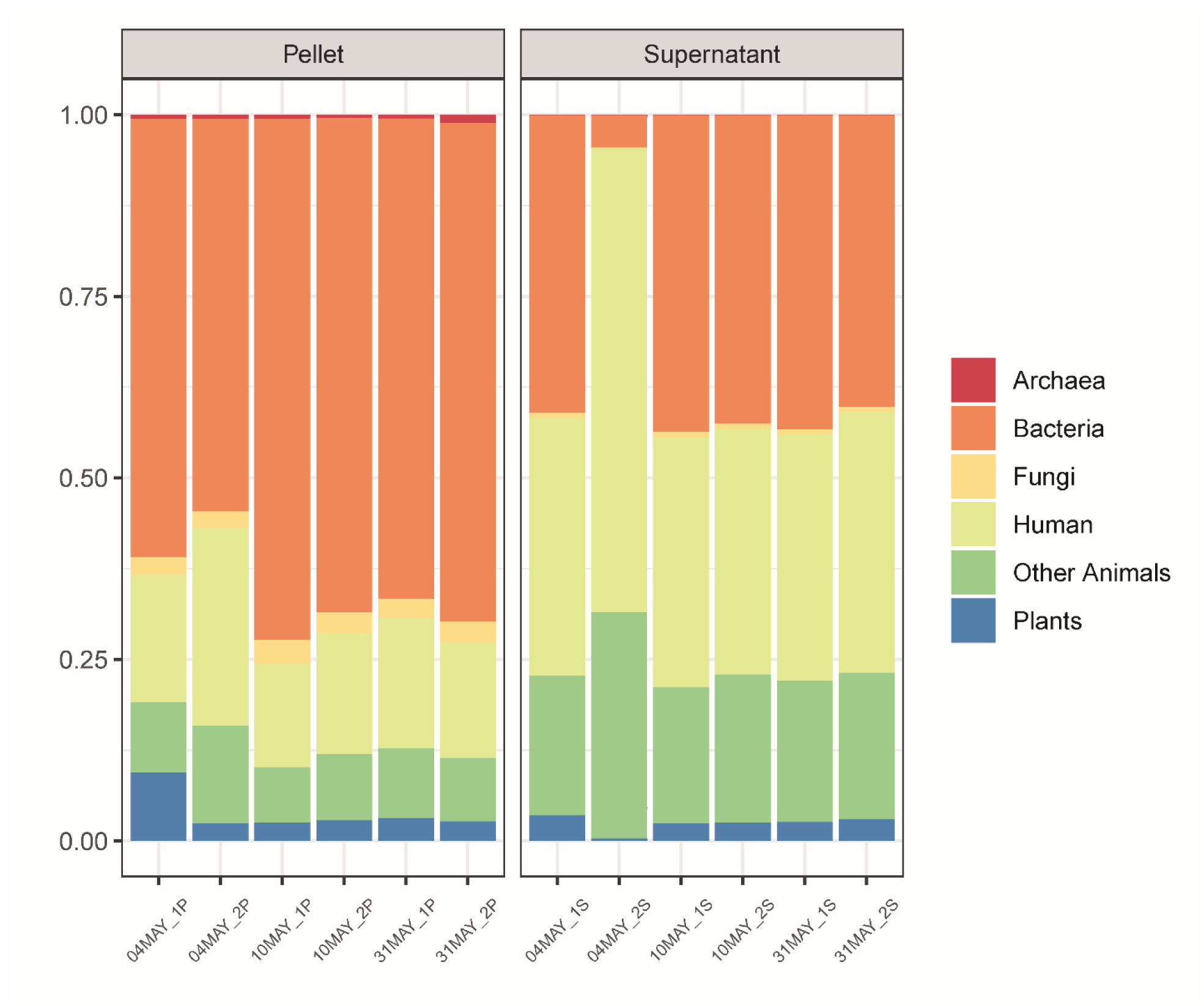
Relative taxonomic (cross-kingdom) distribution in pellet and supernatant fractions from subway air samples (MetaSUB method). Relative taxonomic (cross-kingdom) distribution for subway air samples (N=6) where the intermediate pellet (N=6) and supernatant (N=6) fractions were processed separately with the MetaSUB method.

The random forest classification analysis, where species-level features were scored by their ability to correctly classify the pellet and supernatant groups, had a perfect out-of-bag error of 0%, and the permutation test was statistically significant (*P*>0.001). In the entire dataset, 6.0% of the features were assigned as fungi and 0.3% were assigned as archaea, while among the 100 species with the highest variable importance in our classification model, 56 were fungi and two where archaea. Among the top 50 species, 30 were fungi and one archaea. The top 30 most important features are shown in Figure 13.

**Figure 13.**
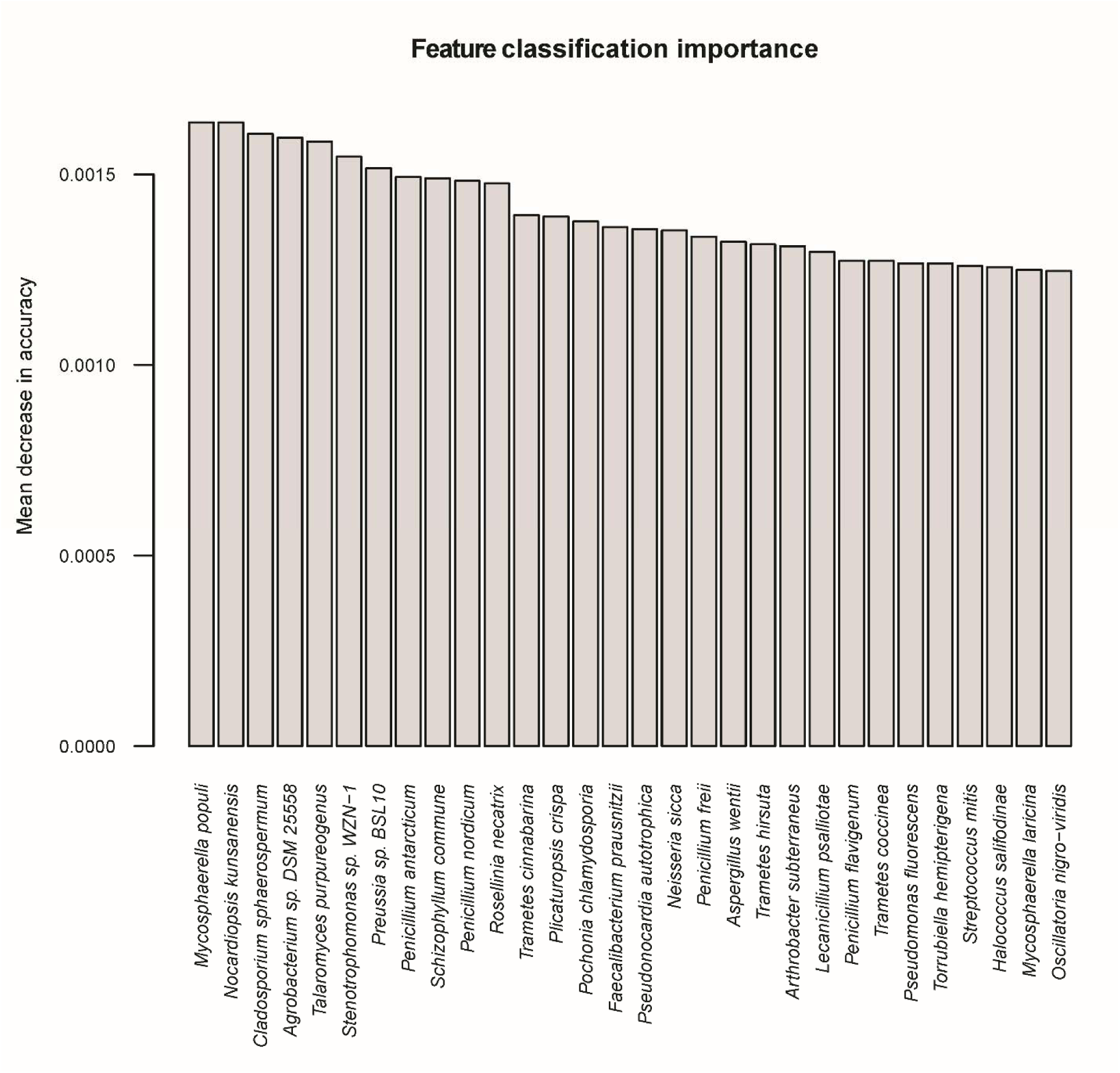
Random forest classification analysis on pellet and supernatant fractions from subway air samples (MetaSUB method). Random forest classification analysis of subway air samples (N=6) where the intermediate pellet (N=6) and supernatant (N=6) fractions were processed separately with the MetaSUB method, showing taxonomic features with the highest classification variable importance for correctly identifying the pellet and supernatant fractions.

## DISCUSSION

Here, we have demonstrated a new custom, multi-component DNA isolation method (“the MetaSUB method”) optimized for SMS-based aerosol microbiome research. By processing the entire filter extract, in combination with thorough chemical, enzymatic and mechanical lysis and DNA purification using magentic beads, the MetaSUB method drastically improves the DNA yield from low biomass air samples and reduces the risk of introducing microbiome profile bias. Comprehensive performance benchmarking of the MetaSUB method against two other state-of-the-art DNA isolation methods was done with both a mock microbial community and real-world subway air samples. The benchmarking revealed that the MetaSUB method obtains significantly higher DNA yields from subway air samples than the other two methods, which is an important performance parameter for successful implementation of SMS on low biomass air samples. SMS of subway air samples revealed that the MetaSUB method resulted in higher numbers of assigned reads than the other two methods, reported higher diversity than Zymobiomics, and gave better representation of certain fungal species than Jiang. All three DNA isolation methods performed similarly well on mock microbial community samples, both in terms of DNA yield and community representation. As part of this study, we have also described an end-to-end air sampling, filter processing and DNA isolation method (“the end-to-end MetaSUB method”) optimized for SMS-based aerosol microbiome research. The end-to-end MetaSUB method relies on the use of SASS 3100 high-volume electret microfibrous filter-based air samplers and was shown to be capable of recovering sufficient DNA yields from short-duration subway air samples, which corresponded to ∼8 m^3^ of air sampled (30 minutes sampling at 265 LPM) in this study, to facilitate high temporal resolution SMS-based aerosol microbiome investigations.

The performence evaluation of the three DNA isolation methods (MetaSUB, Jiang and Zymobiomics) revealed no significant differences regarding total DNA and 16S rRNA gene copy yields when isolating DNA from mock microbial community samples (Figure 2). Furthermore, SMS of mock community samples showed that the three methods gave highly similar representation of the ten microbial species present in the mock community (Figure 3). However, on subway air samples, the MetaSUB method outperformed both Jiang and Zymobiomics regarding total DNA and 16S rRNA gene copy yields (Figure 4). SMS analyses of subway air samples that had been split and isolated with either MetaSUB and Jiang or MetaSUB and Zymobiomics revealed significant differences among the three methods. The numbers of assigned reads were higher for MetaSUB in both comparisons, which is congruent with the higher DNA yields seen for the MetaSUB method (Figure 4). We also observed significantly higher α-diversity estimates for MetaSUB compared to Zymobiomics (Figure 6). One of the three samples processed with Jiang showed higher α-diversity than all three MetaSUB samples, while the other two Jiang samples showed substantially lower diversity estimates (Figure 6), which rendered the comparison against MetaSUB non-significant for all α-diversity indices. We have no conclusive explanation for this pattern; however, we observed that the two low-scoring Jiang samples had high duplicate sequence read proportions (62.4% and 71.8%) compared to all other samples (average: 18.6%), and postulate that the variable performance may be related to the recovery of insufficient DNA yields from two of the Jiang samples to allow for reliable SMS. Furthermore, the random forest classification analysis indicates that the Jiang method does not produce the same representation for certain fungal species as the MetaSUB method, since out of the 100 most important species for distinguishing between MetaSUB and Jiang processed samples, 20 where fungal, while across the entire dataset, only 6% of the species were fungal. All of these 20 fungal species had higher representation in MetaSUB samples (Figure S4).

Our findings highlight the importance of benchmarking DNA isolation methods with both mock communities and real-world samples since the complexity found in the real-world environment is not easily recreated. The observed DNA yield differences among the three methods can probably be attributed to a combination of sub-process efficiency differences, since the methods rely on different combinations of lysis (chemical, enzymatic, and/or mechanical), inhibitor removal and sample clean-up, and DNA purification (magnetic beads and silica spin filters) principles (Table 1). During customization of DNA isolation methods it is therefore important to keep in mind that even subtle procedural differences, including choice of bead solution, intensity and time settings for the bead beating process [47, 48], and different enzyme combinations, may have a large effect on the ultimate biomass lysis efficiency [31]. By replacing the spin columns in the PowerSoil Kit with AMPure XP Beads (magnetic bead purification), Jiang et al. [28] observed a three-fold increase in DNA yield. The multi-component MetaSUB method was developed by adopting and customizing sub-processes from several different DNA isolation methods in an effort to ensure maximized DNA recovery and thorough unbiased biomass lysis. Note that for the performance benchmarking of DNA isolation methods in this study, only the intermediate pellet fraction of the MetaSUB method was used to facilitate an equal comparison between the three different DNA isolation methods (Figure 1). The intermediate supernatant fraction would normally also be included in the MetaSUB method and would have constituted approximately 72% of additional DNA, thereby making the DNA yield differences even more pronounced.

Since the filter extraction procedure in the MetaSUB method produces intermediate pellet and supernatant fraction that are combined before DNA purification, we investigated differences in DNA amount and composition (diversity) between the two fractions in an effort to better understand the benefit of including supernatants (i.e., increased DNA yield) and the risk of not including them (i.e., microbiome profile bias). The observed microbial diversity in paired pellet and supernatant samples was highly similar at the phylum (Table 5), family (Table 5; Figure 9), genus (Table 5) and species (Table 5) levels. Note, however, with direct examination of only the most abundant taxonomic groups in Table 5 and Figure 9, the similarities do not necessarily extend to groups with low abundance. While we did not find any differences among the pellet and supernatant samples in α-diversity (Figure 10), which describes within-sample diversity, there was significant diversity nested among samples, of which the pellet/supernatant grouping explained 51.7% (Figure 11). The cross-kingdom analyses revealed differences in the taxonomic composition of pellet and supernatant samples (Figure 12). While human DNA constituted a relatively large proportion of eukaryotic reads, it did not account for all of the difference observed among pellet and supernatant samples within this kingdom; on average, human reads constituted 18% of assigned reads in pellets and 42% in supernatants based on the cross-kingdom analysis (Figure 12). Human reads reported by One Codex also had a higher relative abundance in supernatants (31% and 67% of assigned reads in pellets and supernatants, respectively). Features assigned as archaea were exclusively observed in pellets; however, caution should be used when interpreting these results, since only eleven features were assigned to this kingdom. The random forest classification model revealed that fungi were particularly important in separating pellet and supernatant samples, especially when accounting for the relatively low representation of fungi across all samples. A recent study by Mbareche et al. has shown that the use of traditional processing methods, e.g., filter extract processing where the supernatant fraction is discarded after a centrifugation step, may lead to an underrepresentation of fungi [49]. In conclusion, concerning the most abundant microbial groups and within-sample diversity estimates, there is little difference between the pellet and supernatant fractions. However, the between-sample diversity analyses show that potentially important diversity may be lost if the entire filter extract is not processed, and that an appreciable amount of this diversity is nested in fungi. In addition, a more general but potentially important reason for processing the entire filter extract in the context of high-volume filter-collected air samples is the variable resistance different types of microorganisms have against sampling-associated stress factors. While stress-resistant microorganisms may be relatively unaffected by sampling-associated stress, stress-sensitive organisms, e.g. Gram-negative bacteria, may become membrane-impaired, ruptured or even completely lysed due to sampling-associated desiccation during high-volume dry filter collection and subsequent osmotic shock during liquid filter extraction. DNA that becomes liberated from membrane-impaired, ruptured or lysed microorganisms will generally not be recovered by standard centrifugation or filtration processes intended for intact organism capture, and may therefore remain in the supernatant or filtrate fraction.

Taken together, the demonstrated performance of the MetaSUB method, including drastically improved DNA yield from subway air samples and reduced risk of microbiome profile bias, highlights the benefit of isolating DNA from the entire filter extract. However, the need for isolating DNA from a relatively large sample volume, a 10 ml filter extract in this work, limits the available selection of out-of-the-box commercial DNA isolation kits and introduces a customization need to ensure reliable performance regarding thorough unbiased biomass lysis, sufficient inhibitor removal and sample clean-up, and efficient DNA recovery. The custom, multi-component MetaSUB method is therefore a relatively hands-on (manual), labor-intensive DNA isolation method compared to many out-of-the-box commercial DNA isolation kits. However, an experienced operator can perform the MetaSUB method, including all processing and incubation steps, in approximately three hours, while the estimated total processing time for 12 air samples is approximately four hours. Furthermore, even without considering the associated benefits of isolating DNA from the entire filter extract, the use of a custom, multi-component DNA isolation method, including extensively modified commercial DNA isolation kits, appears to be necessary to overcome the unique and inherent challenges associated with SMS-based aerosol microbiome research in complex low biomass air environments [2, 21, 28, 29].

## CONCLUSIONS

By demonstrating and benchmarking a new custom, multi-component DNA isolation method (the MetaSUB method) optimized for SMS-based aerosol microbiome research, this study contributes to improved selection, harmonization, and standardization of DNA isolation methods. In the context of SMS-based aerosol microbiome research in low biomass air environments, our findings highlight the importance of ensuring end-to-end sample integrity and using DNA isolation methods with well-defined performance characteristics regarding both DNA yield and community representation. A comprehensive performance benchmarking of the MetaSUB method against two other state-of-the-art DNA isolation methods (Jiang and Zymobiomics) was done with both a mock microbial community and real-world subway air samples. All three DNA isolation methods performed similarly well on mock community samples, both in terms of DNA yield and community representation. However, the MetaSUB method obtained significantly higher DNA yields than the other two methods from subway air samples, which is an important performance parameter for successful implementation of SMS on low biomass air samples. We also observed significant differences regarding SMS-based community representation across the three methods when applying them to subway air samples. The MetaSUB method reported higher α-diversity estimates than Zymobiomics, while Jiang appeared to underrepresent certain fungal species. By processing the entire filter extract, in combination with thorough chemical, enzymatic and mechanical biomass lysis, and efficient DNA recovery using magnetic beads, the MetaSUB method may drastically improve the DNA yield from low biomass air samples and reduce the risk of aerosol microbiome profile bias. Taken together, the demonstrated performance characteristics suggest the MetaSUB method could be used to improve the quality of SMS-based aerosol microbiome research in low biomass air environments. Furthermore, the MetaSUB method, when used in combination with the described high-volume filter-based air sampling, filter processing and DNA isolation scheme (the end-to-end MetaSUB method), could be used to improve the temporal resolution in aerosol microbiome research by reducing the sampling time required to obtain sufficient DNA yields for SMS analysis.

## DECLARATIONS

### Ethics approval and consent to participate

Not applicable

### Consent for publication

Not applicable

### Availability of data and material

The sequence data has been deposited in the NCBI Sequence Read Archive under Bioproject ID# PRJNA542423 (https://www.ncbi.nlm.nih.gov/bioproject/PRJNA542423).

### Authors’ contributions

MD conceived, designed and led the study. JG performed the data analysis. LV-M contributed to the experimental work and the data analysis. KO-B performed the experimental work and contributed to the data analysis. All authors contributed to the manuscript writing and approved the final manuscript.

## Acknowledgements

The sequencing service was provided by the Norwegian Sequencing Centre (www.sequencing.uio.no), a national technology platform hosted by the University of Oslo and supported by the “Functional Genomics” and “Infrastructure” programs of the Research Council of Norway and the Southeastern Regional Health Authorities.

## SUPPLEMENTAL ONLINE MATERIAL

Figure S1 – Rarefaction curves with α-diversity measures: “Observed”, “Shannon”, and “Simpson” for subway air samples (N=6) that were split and processed with the MetaSUB (N=3) and Jiang (N=3) or MetaSUB (N=3) and Zymobiomics (N=3) methods.

Figure S2 – Rarefaction curves with α-diversity measures: “Observed”, “Shannon”, and “Simpson” for the intermediate pellet (N=6) and supernatant (N=6) fractions from subway air samples (N=6) processed separately with the MetaSUB method.

Figure S3 – Proportion of total DNA and 16S rRNA gene copy yield found in the supernatant fractions, referencing the total yield in the combined pellet and supernatant fractions, from subway air samples (N=24) where the intermediate pellet and supernatant fractions were processed separately with the MetaSUB method.

Figure S4 – The 20 fungal species that were among the top 100 species from the random forest classification analysis of subway air samples (N=3) that were split and processed with the MetaSUB (N=3) and Jiang (N=3) methods, where *Z*-score distributions were compared with linear models.

## REFERENCES

1. Yao M: Bioaerosol: A bridge and opportunity for many scientific research fields. Journal of Aerosol Science 2018, 115:108–112, http://dx.doi.org/10.1016/j.jaerosci.2017.07.010.

2. King P, Pham LK, Waltz S, Sphar D, Yamamoto RT, Conrad D, Taplitz R, Torriani F, Forsyth RA: Longitudinal Metagenomic Analysis of Hospital Air Identifies Clinically Relevant Microbes. Plos One 2016, 11(8):e0160124, http://dx.doi.org/10.1371/journal.pone.0160124.

3. Choi JY, Zemke J, Philo SE, Bailey ES, Yondon M, Gray GC: Aerosol Sampling in a Hospital Emergency Room Setting: A Complementary Surveillance Method for the Detection of Respiratory Viruses. Frontiers in Public Health 2018, 6:174–174, http://dx.doi.org/10.3389/fpubh.2018.00174.

4. Nguyen TT, Poh MK, Low J, Kalimuddin S, Thoon KC, Ng WC, Anderson BD, Gray GC: Bioaerosol Sampling in Clinical Settings: A Promising, Noninvasive Approach for Detecting Respiratory Viruses. Open forum infectious diseases 2017, 4(1):ofw259, http://dx.doi.org/10.1093/ofid/ofw259.

5. Prost K, Kloeze H, Mukhi S, Bozek K, Poljak Z, Mubareka S: Bioaerosol and surface sampling for the surveillance of influenza A virus in swine. Transboundary and emerging diseases 2019, http://dx.doi.org/10.1111/tbed.13139.

6. Osman S, La Duc MT, Dekas A, Newcombe D, Venkateswaran K: Microbial burden and diversity of commercial airline cabin air during short and long durations of travel. International Society for Microbial Ecology (ISME) Journal 2008, 2:482–497, http://dx.doi.org/10.1038/ismej.2008.11.

7. Weiss H, Hertzberg VS, Dupont C, Espinoza JL, Levy S, Nelson K, Norris S, Team TFR: The Airplane Cabin Microbiome. Microbial Ecology 2019, 77(1):87–95, http://dx.doi.org/10.1007/s00248-018-1191-3.

8. Zanni S, Lalli F, Foschi E, Bonoli A, Mantecchini L: Indoor Air Quality Real-Time Monitoring in Airport Terminal Areas: An Opportunity for Sustainable Management of Micro-Climatic Parameters. Sensors 2018, 18(11):3798, http://dx.doi.org/10.3390/s18113798.

9. Wagar E: Bioterrorism and the Role of the Clinical Microbiology Laboratory. Clinical Microbiology Reviews 2016, 29(1):175–189, http://dx.doi.org/10.1128/CMR.00033-15.

10. Amann RI, Ludwig W, Schleifer K-H: Phylogenetic identification and in situ detection of individual microbial cells without cultivation. Microbiology and Molecular Biology Reviews 1995, 59(1):143–169,

11. Radosevich JL, Wilson WJ, Shinn JH, DeSantis TZ, Andersen GL: Development of a high-volume aerosol collection system for the identification of air-borne micro-organisms. Lett Appl Microbiol 2002, 34(3):162–167, http://dx.doi.org/10.1046/j.1472-765x.2002.01048.x.

12. Toivola M, Alm S, Reponen T, Kolari S, Nevalainen A: Personal exposures and microenvironmental concentrations of particles and bioaerosols. J Environ Monit 2002, 4(1):166–174,

13. Eduard W, Heederik D: Methods for quantitative assessment of airborne levels of noninfectious microorganisms in highly contaminated work environments. American Industrial Hygiene Association Journal 1998, 59(2):113–127, http://dx.doi.org/10.1080/15428119891010370.

14. Leung MH, Wilkins D, Li EK, Kong FK, Lee PK: Indoor-air microbiome in an urban subway network: diversity and dynamics. Appl Environ Microb 2014, 80(21):6760–6770, http://dx.doi.org/10.1128/AEM.02244-14.

15. Triado-Margarit X, Veillette M, Duchaine C, Talbot M, Amato F, Minguillon MC, Martins V, de Miguel E, Casamayor EO, Moreno T: Bioaerosols in the Barcelona subway system. Indoor Air 2017, 27(3):564–575, http://dx.doi.org/10.1111/ina.12343.

16. Cáliz J, Triadó-Margarit X, Camarero L, Casamayor EO: A long-term survey unveils strong seasonal patterns in the airborne microbiome coupled to general and regional atmospheric circulations. Proceedings of the National Academy of Sciences 2018, 115(48):12229–12234, http://dx.doi.org/10.1073/pnas.1812826115.

17. Hanson B, Zhou Y, Bautista EJ, Urch B, Speck M, Silverman F, Muilenberg M, Phipatanakul W, Weinstock G, Sodergren E et al: Characterization of the bacterial and fungal microbiome in indoor dust and outdoor air samples: a pilot study. Environmental Science: Processes & Impacts 2016, 18(6):713–724, http://dx.doi.org/10.1039/c5em00639b.

18. The Human Microbiome Project Consortium: Structure, function and diversity of the healthy human microbiome. Nature 2012, 486:207–214, http://dx.doi.org/10.1038/nature11234.

19. Afshinnekoo E, Meydan C, Chowdhury S, Jaroudi D, Boyer C, Bernstein N, Maritz JM, Reeves D, Gandara J, Chhangawala S et al: Geospatial Resolution of Human and Bacterial Diversity with City-Scale Metagenomics. Cell Systems 2015, 1(1):72–87, http://dx.doi.org/10.1016/j.cels.2015.01.001.

20. Biller SJ, Berube PM, Dooley K, Williams M, Satinsky BM, Hackl T, Hogle SL, Coe A, Bergauer K, Bouman HA et al: Marine microbial metagenomes sampled across space and time. Scientific data 2018, 5:180176, http://dx.doi.org/10.1038/sdata.2018.176.

21. Yooseph S, Andrews-Pfannkoch C, Tenney A, McQuaid J, Williamson S, Thiagarajan M, Brami D, Zeigler-Allen L, Hoffman J, Goll JB et al: A Metagenomic Framework for the Study of Airborne Microbial Communities. Plos One 2013, 8(12):e81862, http://dx.doi.org/10.1371/journal.pone.0081862.

22. Cao C, Jiang W, Wang B, Fang J, Lang J, Tian G, Jiang J, Zhu TF: Inhalable microorganisms in Beijing’s PM2.5 and PM10 pollutants during a severe smog event. Environmental Science & Technology 2014, 48(3):1499–1507, http://dx.doi.org/10.1021/es4048472.

23. Eisenhofer R, Minich JJ, Marotz C, Cooper A, Knight R, Weyrich LS: Contamination in Low Microbial Biomass Microbiome Studies: Issues and Recommendations. Trends Microbiol 2019, 27(2):105–117, http://dx.doi.org/https://doi.org/10.1016/j.tim.2018.11.003.

24. Dybwad M, Skogan G, Blatny JM: Comparative Testing and Evaluation of Nine Different Air Samplers: End-to-End Sampling Efficiencies as Specific Performance Measurements for Bioaerosol Applications. Aerosol Science and Technology 2014, 48(3):282–295, http://dx.doi.org/10.1080/02786826.2013.871501.

25. Behzad H, Gojobori T, Mineta K: Challenges and opportunities of airborne metagenomics. Genome Biology and Evolution 2015, 7(5):1216–1226, http://dx.doi.org/10.1093/gbe/evv064.

26. Tringe SG, Hugenholtz P: A renaissance for the pioneering 16S rRNA gene. Current opinion in microbiology 2008, 11(5):442–446, http://dx.doi.org/10.1016/j.mib.2008.09.011.

27. Luhung I, Wu Y, Ng CK, Miller D, Cao B, Chang VW: Protocol Improvements for Low Concentration DNA-Based Bioaerosol Sampling and Analysis. Plos One 2015, 10(11):e0141158, http://dx.doi.org/10.1371/journal.pone.0141158.

28. Jiang W, Liang P, Wang B, Fang J, Lang J, Tian G, Jiang J, Zhu TF: Optimized DNA extraction and metagenomic sequencing of airborne microbial communities. Nature Protocols 2015, 10:768, http://dx.doi.org/10.1038/nprot.2015.046.

29. Dommergue A, Amato P, Tignat-Perrier R, Magand O, Thollot A, Joly M, Bouvier L, Sellegri K, Vogel T, Sonke JE et al: Methods to Investigate the Global Atmospheric Microbiome. Frontiers in Microbiology 2019, 10:00243, http://dx.doi.org/10.3389/fmicb.2019.00243.

30. Tighe S, Afshinnekoo E, Rock TM, McGrath K, Alexander N, McIntyre A, Ahsanuddin S, Bezdan D, Green SJ, Joye S et al: Genomic Methods and Microbiological Technologies for Profiling Novel and Extreme Environments for the Extreme Microbiome Project (XMP). Journal of Biomolecular Techniques 2017, 28(1):31–39, http://dx.doi.org/10.7171/jbt.17-2801-004.

31. Yuan S, Cohen DB, Ravel J, Abdo Z, Forney LJ: Evaluation of methods for the extraction and purification of DNA from the human microbiome. Plos One 2012, 7(3):e33865, http://dx.doi.org/10.1371/journal.pone.0033865.

32. Abusleme L, Hong BY, Dupuy AK, Strausbaugh LD, Diaz PI: Influence of DNA extraction on oral microbial profiles obtained via 16S rRNA gene sequencing. Journal of Oral Microbiology 2014, 6:23990, http://dx.doi.org/10.3402/jom.v6.23990.

33. Mbareche H, Brisebois E, Veillette M, Duchaine C: Bioaerosol sampling and detection methods based on molecular approaches: No pain no gain. Science of The Total Environment 2017, 599–600:2095–2104, http://dx.doi.org/10.1016/j.scitotenv.2017.05.076.

34. The Human Microbiome Project Consortium: A framework for human microbiome research. Nature 2012, 486:215–221, http://dx.doi.org/10.1038/nature11209.

35. Gilbert JA, Jansson JK, Knight R: Earth Microbiome Project and Global Systems Biology. mSystems 2018, 3(3):e00217, http://dx.doi.org/10.1128/mSystems.00217-17.

36. Kopf A, Bicak M, Kottmann R, Schnetzer J, Kostadinov I, Lehmann K, Fernandez-Guerra A, Jeanthon C, Rahav E, Ullrich M et al: The ocean sampling day consortium. GigaScience 2015, 4(1):s13742, http://dx.doi.org/10.1186/s13742-015-0066-5.

37. Lear G, Dickie I, Banks J, Boyer S, Buckley HL, Buckley TR, Cruickshank R, Dopheide A, Handley KM, Hermans S et al: Methods for the extraction, storage, amplification and sequencing of DNA from environmental samples. New Zealand Journal of Ecology 2018, 42(1):10–50, http://dx.doi.org/10.20417/nzjecol.42.9.

38. Al-Hebshi NN, Baraniya D, Chen T, Hill J, Puri S, Tellez M, Hasan NA, Colwell RR, Ismail A: Metagenome sequencing-based strain-level and functional characterization of supragingival microbiome associated with dental caries in children. Journal of Oral Microbiology 2018, 11(1):1557986, http://dx.doi.org/10.1080/20002297.2018.1557986.

39. Honeyman AS, Day ML, Spear JR: Regional fresh snowfall microbiology and chemistry are driven by geography in storm-tracked events, Colorado, USA. PeerJ 2018, 6:e5961, http://dx.doi.org/10.7717/peerj.5961.

40. Zaikova E, Goerlitz DS, Tighe SW, Wagner NY, Bai Y, Hall BL, Bevilacqua JG, Weng MM, Samuels-Fair MD, Johnson SS: Antarctic Relic Microbial Mat Community Revealed by Metagenomics and Metatranscriptomics. Frontiers in Ecology and Evolution 2019, 7(1):10.3389, http://dx.doi.org/10.3389/fevo.2019.00001.

41. Trivedi CB, Lau GE, Grasby SE, Templeton AS, Spear JR: Low-Temperature Sulfidic-Ice Microbial Communities, Borup Fiord Pass, Canadian High Arctic. Frontiers in Microbiology 2018, 9:01622, http://dx.doi.org/10.3389/fmicb.2018.01622.

42. Liu CM, Aziz M, Kachur S, Hsueh P-R, Huang Y-T, Keim P, Price LB: BactQuant: an enhanced broad-coverage bacterial quantitative real-time PCR assay. BMC Microbiol 2012, 12:56–56, http://dx.doi.org/10.1186/1471-2180-12-56.

43. Minot SS, Krumm N, Greenfield NB: One Codex: A Sensitive and Accurate Data Platform for Genomic Microbial Identification. bioRxiv 2015:027607, http://dx.doi.org/10.1101/027607.

44. McMurdie PJ, Holmes S: phyloseq: an R package for reproducible interactive analysis and graphics of microbiome census data. PloS one 2013, 8(4):e61217,

45. Huson DH, Auch AF, Qi J, Schuster SC: MEGAN analysis of metagenomic data. Genome Res 2007, 17(3):377–386,

46. Liaw A, Wiener M: Classification and regression by randomForest. R news 2002, 2(3):18–22,

47. Miller DN, Bryant JE, Madsen EL, Ghiorse WC: Evaluation and optimization of DNA extraction and purification procedures for soil and sediment samples. Appl Environ Microb 1999, 65(11):4715–4724,

48. Griffiths LJ, Anyim M, Doffman SR, Wilks M, Millar MR, Agrawal SG: Comparison of DNA extraction methods for Aspergillus fumigatus using real-time PCR. Journal of Medical Microbiology 2006, 55(9):1187–1191, http://dx.doi.org/10.1099/jmm.0.46510-0.

49. Mbareche H, Veillette M, Teertstra W, Kegel W, Bilodeau GJ, Wösten HAB, Duchaine C: Fungal Cells Recovery From Air Samples: A Tale of Loss And Gain. Appl Environ Microb 2019:02941, http://dx.doi.org/10.1128/aem.02941-18.

