## Supplemental Figures for "Performance evaluation of a new custom, multi-component DNA isolation method optimized for use in shotgun metagenomic sequencing-based aerosol microbiome research"

**Figure S1** – Rarefaction curves with α-diversity measures: "Observed", "Shannon", and "Simpson" for subway air samples (N=6) that were split and processed with the MetaSUB (N=3) and Jiang (N=3) or MetaSUB (N=3) and Zymobiomics (N=3) methods.

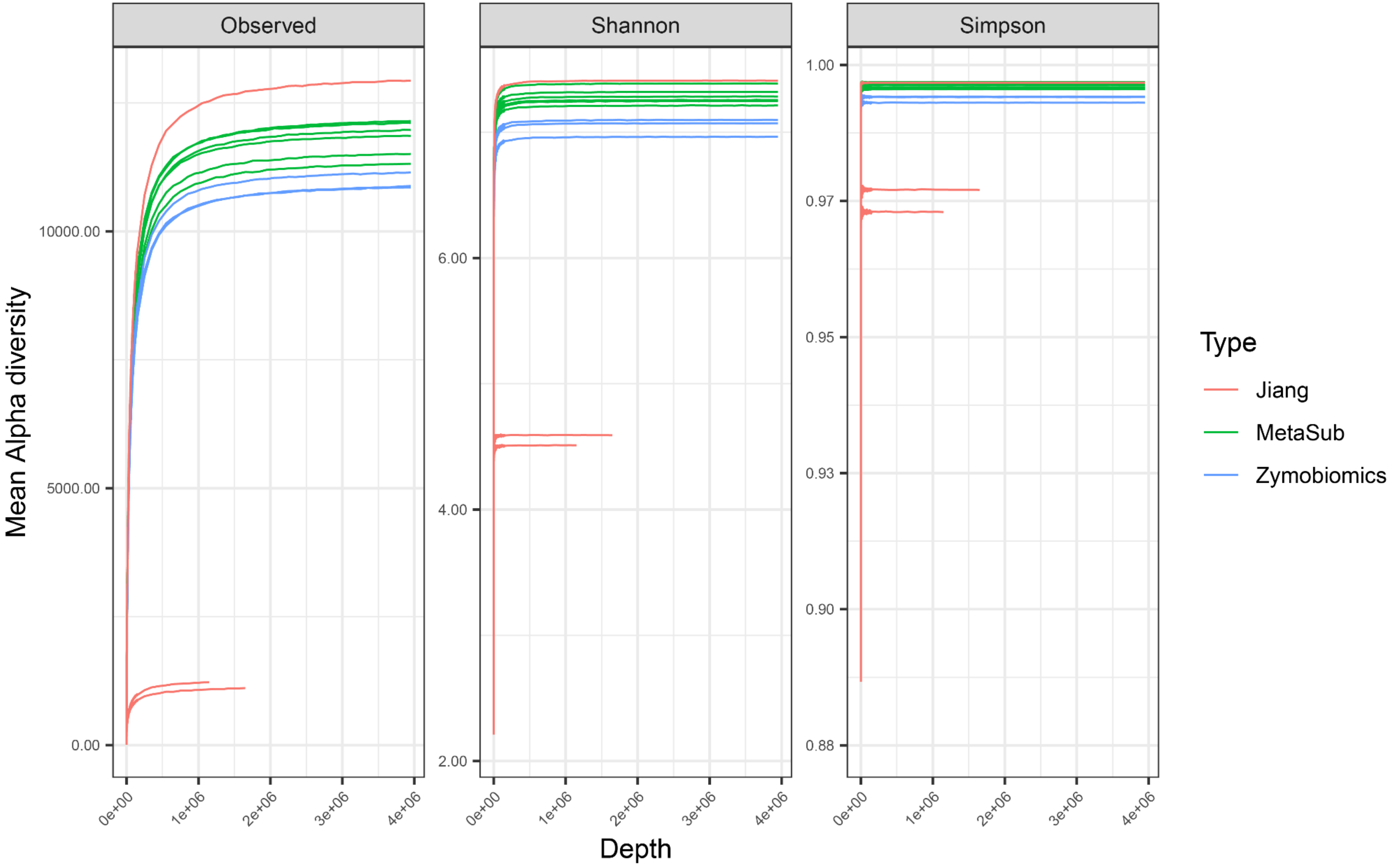

**Figure S2** – Rarefaction curves with α-diversity measures: "Observed", "Shannon", and "Simpson" for the intermediate pellet (N=6) and supernatant (N=6) fractions from subway air samples (N=6) processed separately with the MetaSUB method.

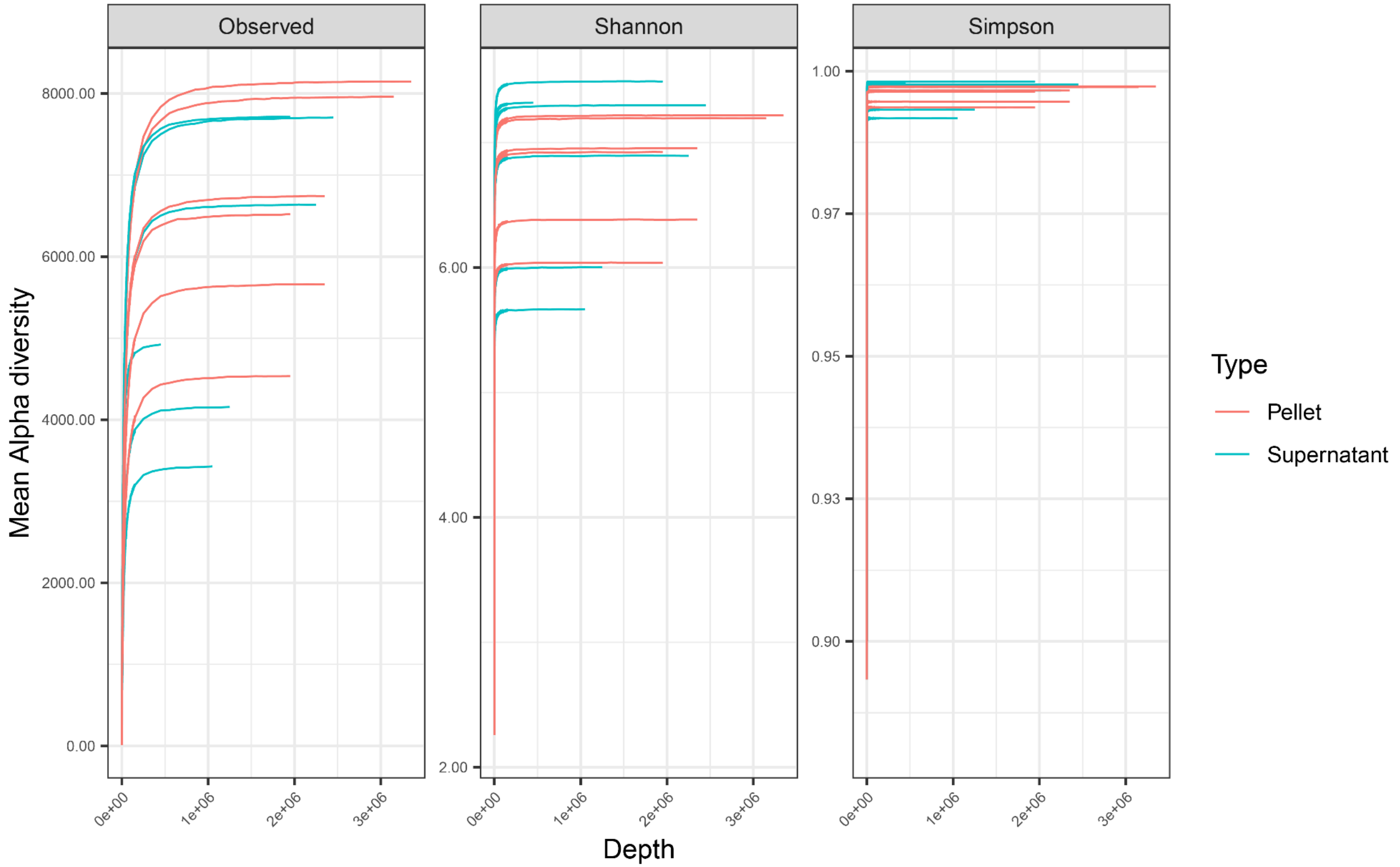

**Figure S3** – Proportion of total DNA and 16S rRNA gene copy yield found in the supernatant fractions, referencing the total yield in the combined pellet and supernatant fractions, from subway air samples (N=24) where the intermediate pellet and supernatant fractions were processed separately with the MetaSUB method.

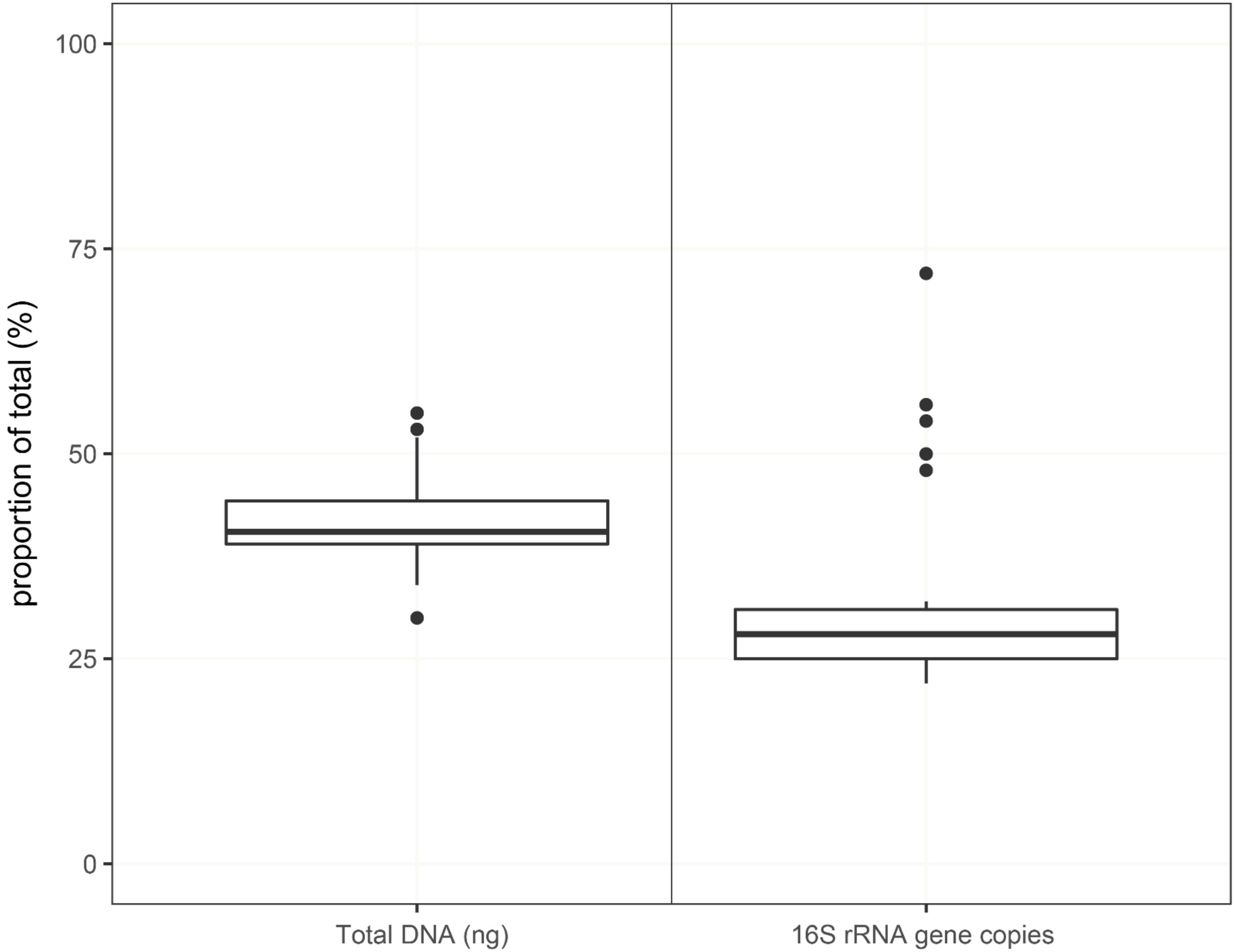

**Figure S4** – The 20 fungal species that were among the top 100 species from the random forest classification analysis of subway air samples (N=3) that were split and processed with the MetaSUB (N=3) and Jiang (N=3) methods, where *Z*-score distributions were compared with linear models.

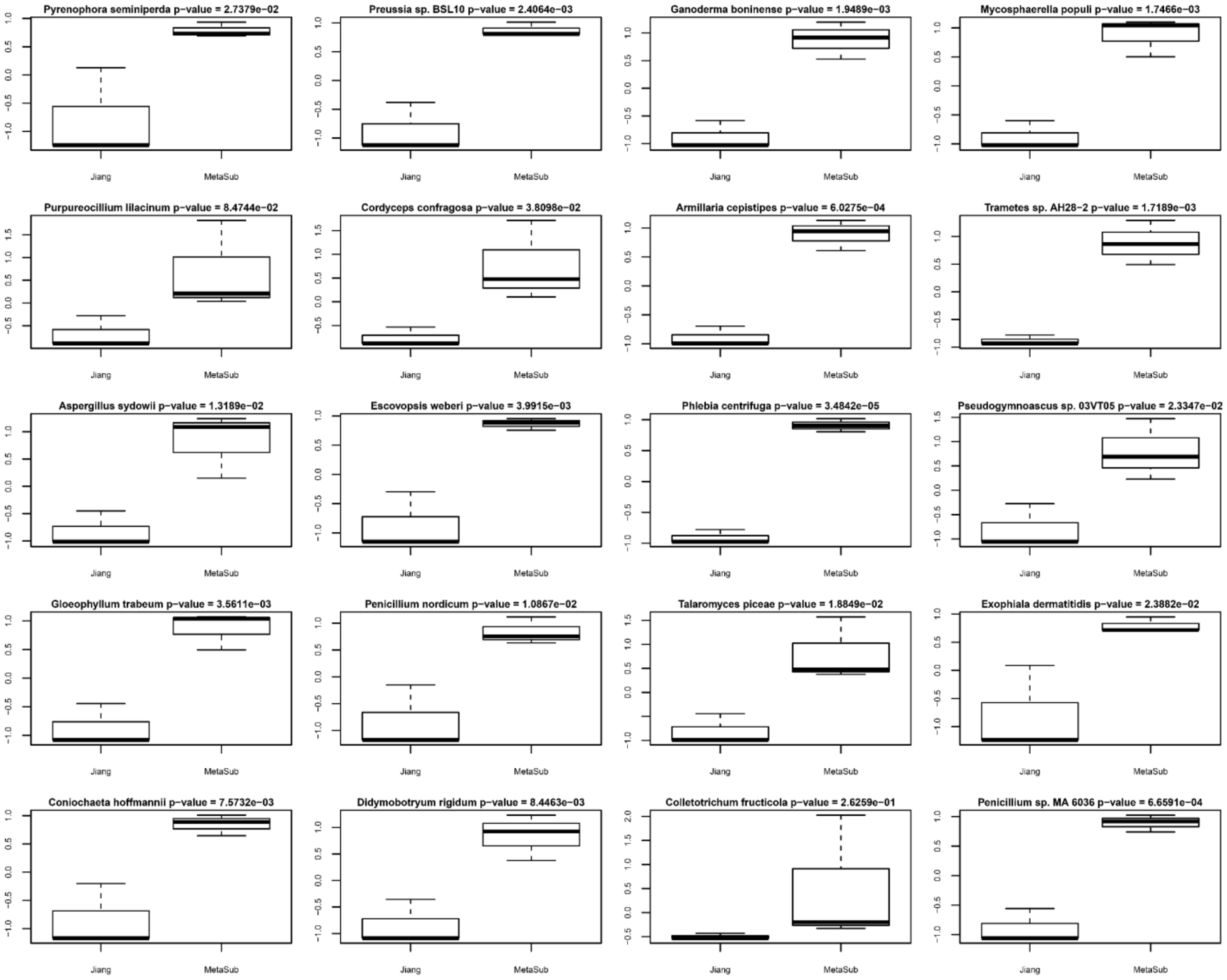
